# A systematic benchmark evaluation of algorithms for 3D tissue alignment and registration with spatial omics

**DOI:** 10.64898/2026.11.23.768628

**Authors:** Jianing Yao, Jan Matthias, Guan Gui, Vani Padmakumar, An Wang, Shu Luan, Yiqun Chen, Laurent Younes, Kristen R. Maynard, Keri Martinowich, Stephanie C. Hicks

## Abstract

Recent advances in spatial omics have enabled tissue atlases spanning serial sections, donors, and experimental technologies. Aligning these datasets is essential for comparing spatial organization, reconstructing three-dimensional tissue architecture, and registering sections to a common coordinate framework. However, systematic comparisons of alignment accuracy, computational efficiency, and usability remain limited. Here, we benchmarked 15 tissue alignment and registration algorithms using experimental and simulated spatial transcriptomics datasets across pairwise alignment, three-dimensional reconstruction, cross-resolution alignment, and atlas registration tasks. We assessed spatial-domain and cell-type correspondence, spatial gene-expression similarity, computational scalability, and usability. Simulated datasets with known spatial coordinates additionally enabled direct evaluation of geometric accuracy and reconstruction stability. No single method performed best across all settings: performance depended on the task, spatial resolution, and tissue characteristics, and strong pairwise alignment did not necessarily translate into reliable three-dimensional reconstruction. Evaluation metrics captured complementary aspects of alignment quality, highlighting the importance of using multiple criteria for comprehensive assessment. These findings provide practical guidance for method selection and identify priorities for improving the accuracy, scalability, and accessibility of spatial omics alignment.

## 1 Introduction

The development of spatial transcriptomics technologies has made it possible to measure mRNA expression directly within its tissue context, improving our ability to study tissue microenvironments and their underlying cellular organization [1, 2]. These technologies broadly fall into two categories: image-based approaches, such as Xenium and MERFISH, which resolve transcripts at single-cell resolution [3], and sequencing-based approaches, such as Visium and Visium HD, which capture transcriptome-wide expression across tissue sections [4, 5].

Recent experiments have collected consecutive or serial tissue slices from the same tissue block, which can be analyzed either as individual two-dimensional sections or integrated into a three-dimensional representation. A common data analysis task is to align these slices, either pairwise between two consecutive sections or jointly across three or more sections to reconstruct a three-dimensional volume.

Meaningful biological interpretation often requires understanding how these slices relate to one another in three-dimensional space. To address this, several computational strategies have been proposed: spatial integration, which combines multiple datasets into a shared representation; spatial alignment, which maps coordinates or matches cells across slices within a common spatial frame; and registration, which places slices into an anatomical reference coordinate system [6, 7]. In practice, alignment often begins with integration to identify cross-slice correspondences, followed by coordinate adjustment into a shared spatial frame. Methods implementing these strategies typically rely on one of several common algorithmic approaches, including optimal transport, graph-based matching, point cloud registration, and deep learning-based feature alignment [8–11]. Together, these enable downstream analyses such as 3D reconstruction, common coordinate framework (CCF) registration for cross-donor comparison, and 3D spatial gradient analysis [12]. Although a growing number of methods have been developed in recent years along with review articles of the methods, their relative suitability for specific tasks remains unclear, and a systematic benchmark comparing their performance across diverse experimental settings has yet to be conducted.

In this paper, we present a systematic benchmark evaluation of spatial alignment and registration methods using both synthetic and experimentally derived datasets. We assess their applicability to 3D tissue alignment through four case studies including pairwise and multi-section alignment and registration into a CCF. We additionally generate a synthetic spatial transcriptomics simulation framework to evaluate alignment performance under controlled conditions, quantify computational resource usage, and examine the influence of different evaluation metrics on method rankings. Combining these results with insights from the synthetic data analyses, we provide practical recommendations for users seeking to align tissue sections and for developers addressing open challenges in alignment and registration for spatial omics.

## 2 Results

### 2.1 Overview of benchmarking spatial alignment and registration methods

Here, we present a systematic benchmark of 15 spatial alignment and registration algorithms, selected from 24 surveyed methods, using experimental and simulated spatial transcriptomics datasets. These data capture tissue organization through a series of two-dimensional sections, each containing gene expression measurements linked to spatial coordinates at the cell, spot, or bin level. Aligning these sections establishes spatial correspondence across their separate coordinate systems, enabling comparisons across sections and reconstruction of three-dimensional tissue architecture (**Figure 1A**). The benchmarked methods were GPSA [13], INSPIRE [14], PASTE [6], PASTE2 [15], SANTO [16], SLAT [17], SPACEL [18], Spateo [19], SPIRAL [20], STaCker [21], STAIR [22], STalign [23], STAligner [24], STitch3D [25], and STUtility [26]. These methods encompass probabilistic spatial warping, optimal transport, morphology-based registration, and graph representation (**Figure 1B**). These categories summarize major methodological principles rather than mutually exclusive classes, as individual workflows may combine multiple approaches.

**Figure 1:**
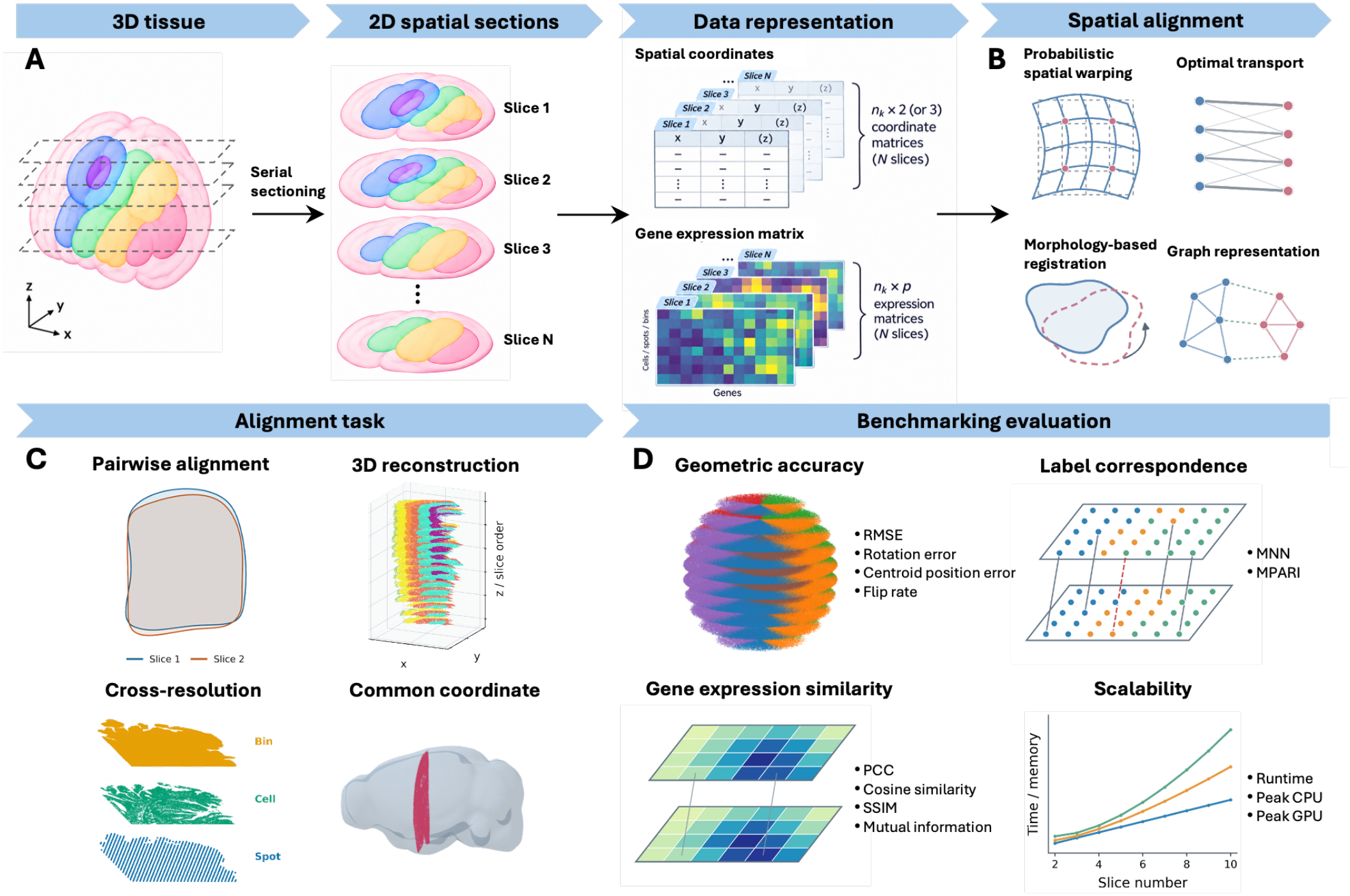
Overview of spatial alignment and registration benchmarking. (**A**) Spatial transcriptomics data representation, from a three-dimensional tissue to serial two-dimensional sections, each represented by spatial coordinates and a gene expression matrix. (**B**) Schematic illustrations of four methodological categories represented among the 15 benchmarked algorithms: probabilistic spatial warping, optimal transport, morphology-based registration, and graph representation. (**C**) Four alignment tasks evaluated using experimental and simulated datasets: pairwise alignment, three-dimensional reconstruction, cross-resolution alignment, and registration to a common coordinate framework. (**D**) Four categories of benchmarking evaluation: geometric accuracy, including root mean squared error (RMSE), rotation error, centroid position error, and flip rate; spatial label correspondence, including mutual nearest-neighbor (MNN) accuracy and mean pairwise adjusted Rand index (MPARI); spatial gene-expression similarity, including Pearson correlation coefficient (PCC), cosine similarity, structural similarity index (SSIM), and mutual information; and computational scalability, including runtime and peak CPU and GPU memory usage as the number of input slices increases.

We considered four tasks: pairwise alignment (i.e., aligning two spatial transcriptomics slices), three-dimensional reconstruction from multiple sections, cross-resolution alignment across experimental platforms, and registration of individual sections to a common coordinate framework (CCF) (**Figure 1C**). Methods were evaluated on the tasks for which applicable workflows and usable outputs were available. Experimental datasets included human dorsolateral prefrontal cortex (DLPFC), mouse hypothalamus, mouse primary motor cortex, and human colon, profiled using Visium, Visium HD, MERFISH, and Xenium, as well as with mouse brain data registered to the Allen CCF for atlas-registration evaluation. Simulated datasets generated using ALBIS [27] provided matched cell-, bin-, and spot-level observations derived from the same underlying simulated tissue and RNA molecules.

Evaluation covered four complementary aspects (**Figure 1D**): geometric accuracy, spatial label correspondence, spatial gene-expression similarity, and computational scalability. Geometric accuracy was assessed using root mean squared error (RMSE), rotation error, centroid position error, and flip rate where the required coordinate references were available. In particular, simulated spatial coordinates enabled direct assessment of alignment error and reconstruction stability, whereas CCF localization was evaluated against published atlas-registered coordinates. Spatial label correspondence was assessed using mutual nearest-neighbor (MNN) accuracy and mean pairwise adjusted Rand index (MPARI). Gene-expression similarity was assessed using Pearson correlation coefficient (PCC), cosine similarity, structural similarity index (SSIM), and mutual information (MI). Computational scalability was characterized by runtime and peak CPU and GPU memory usage as the number of input sections increased, with all experiments conducted under the same hardware resource configuration. We additionally assessed usability based on our experience installing and applying each method. Complementing existing benchmarks [7, 9–11], this study combines multiple alignment tasks, tissues, technologies, and spatial resolutions with controlled simulations, enabling assessment of both biological correspondence and geometric accuracy alongside computational requirements.

### 2.2 Pairwise alignment and three-dimensional reconstruction within donors

Pairwise alignment is a fundamental task in multi-section spatial transcriptomics analysis, supporting both spatial comparisons between sections and subsequent 3D reconstruction. We evaluated pairwise alignment across spot-level Visium and cell-level MERFISH datasets to examine performance across spatial transcriptomics technologies and resolutions. For the 18 within-donor pairs from 12 DLPFC Visium sections, SPACEL and INSPIRE achieved the highest median cortical-layer MNN agreement, exceeding 0.8, with relatively compact score distributions (**Figure 2A**). PASTE2 and STalign also showed high median agreement, while rigid and non-rigid modes of Spateo were competitive but more variable across pairs. STAligner and SANTO had the lowest median scores. Representative overlays illustrated differences in cortical-layer correspondence and revealed a flipped orientation in the PASTE alignment (**Figure 2B**). Gene-expression metrics provided a complementary comparison: SPACEL and INSPIRE had the highest median PCC, whereas method rankings differed somewhat across cosine similarity, SSIM, and MI (**Figure S1A**).

**Figure 2:**
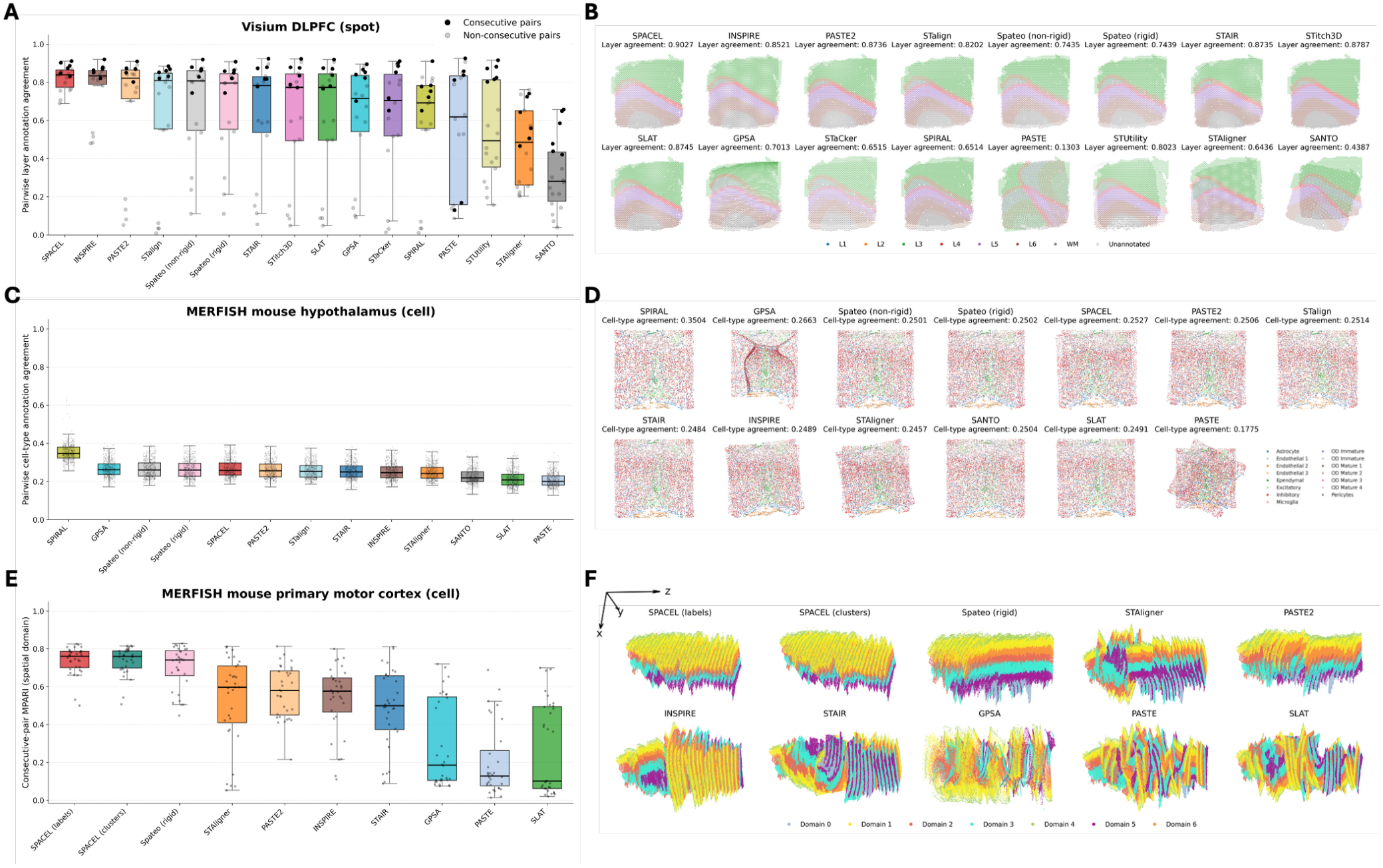
Pairwise alignment and 3D reconstruction performance across spatial transcriptomics datasets. **(A)** Pairwise cortical-layer annotation agreement for the human dorsolateral prefrontal cortex (DLPFC) Visium dataset, evaluated using coordinate-based mutual nearest neighbors (MNN). Each point represents one of 18 within-donor section pairs across three donors; black and gray points indicate consecutive and non-consecutive pairs, respectively. **(B)** Representative aligned overlays of DLPFC sections 151671 and 151672 across methods. Spots are colored by cortical-layer annotation. **(C)** Pairwise cell-type annotation agreement for the mouse hypothalamus MERFISH dataset, evaluated using coordinate-based MNN. Each point represents an available section pair. **(D)** Representative aligned overlays of mouse hypothalamus sections from mouse 3 at depths 0.06 and 0.16. Cells are colored by cell-type annotation. **(E)** Spatial-domain MPARI for 3D reconstructions of the mouse primary motor cortex MERFISH dataset. Each point represents one of 32 consecutive section pairs within the 33-section reconstruction. **(F)** Corresponding 3D reconstructions of the mouse primary motor cortex, with cells colored by spatial domain. In **(A), (C)**, and **(E)**, boxes indicate the interquartile range, center lines indicate the median, and whiskers extend to the most extreme observations within 1.5 times the interquartile range.

Mouse hypothalamus MERFISH data provided a contrasting setting, with single-cell observations, a targeted gene panel, and spatially intermixed cell populations. SPIRAL achieved the highest median cell-type MNN agreement, approximately 0.35, followed by GPSA (**Figure 2C**). However, despite its relatively high score, GPSA produced pronounced non-rigid deformation in the representative alignment, visibly distorting tissue geometry (**Figure 2D**). Its flexible transformation may bring cells with similar molecular profiles closer together without preserving the original tissue shape, illustrating that high cell-type agreement alone does not establish anatomical plausibility. Most remaining methods had median MNN scores near 0.2–0.3, with PASTE and SLAT among the lowest. Gene-expression comparisons likewise placed SPIRAL among the strongest-performing methods, particularly for PCC and SSIM (**Figure S1B**). Cell-type MNN scores were generally lower than cortical-layer MNN scores in DLPFC, partly reflecting the different spatial organization of these labels. Cortical layers form contiguous regions, so cells or spots at corresponding positions in adjacent sections are likely to share a layer label. In contrast, multiple hypothalamic cell types coexist within the same local region. Even when two regions are correctly aligned, neighboring cells across sections may therefore have different cell-type labels and receive a low correspondence score. Thus, lower cell-type MNN scores do not necessarily indicate poorer geometric alignment, and their absolute values should not be directly compared with domain-based scores across datasets. Targeted gene coverage and biological variation between sections may also contribute to the observed differences.

Reconstructing a complete tissue volume introduces an additional challenge: alignment must preserve anatomical continuity across a series of sections, and errors at individual interfaces can propagate through the reconstructed stack. We evaluated this task using the mouse primary motor cortex MERFISH dataset from Zhang et al. [28]. Both label-based and clustering-based SPACEL, and rigid Spateo, achieved the highest median spatial-domain MPARI across consecutive sections (**Figure 2E**). Their reconstructed stacks also displayed comparatively coherent cortical-domain organization (**Figure 2F**). STAligner, PASTE2, INSPIRE, and STAIR showed intermediate median MPARI, whereas GPSA, PASTE, and SLAT had lower medians and more variable section-pair scores. The visual comparisons revealed orientation inconsistencies and tissue distortion in several reconstructions, illustrating errors that are not fully characterized by a single summary score. Gene-expression metrics broadly supported the domain-based comparison, with both SPACEL variants and rigid Spateo showing high PCC and SSIM (**Figure S1C**). The availability of both spatial-domain and cell-type subclass annotations in this dataset allowed us to examine how label choice affects evaluation within the same reconstructed tissue. Subclass-based MPARI scores were consistently lower than domain-based scores, while the overall ranking of methods remained largely consistent (**Figure S2**). Although subclass labels provide a finer-grained description than spatial domains, they retain spatial structure in this tissue. This shared anatomical organization may explain why the relative ranking of methods was broadly consistent despite differences in absolute scores. These results highlight the importance of evaluating reconstructed stacks directly, using complementary spatial and gene-expression metrics, and distinguishing regional tissue correspondence from individual cell-type correspondence.

### 2.3 Cross-technology alignment of a shared tissue region

To evaluate alignment across technologies with differing spatial resolutions and gene coverage, we analyzed tissue sections from a single human colon cancer donor profiled using Visium, Visium HD, and Xenium [4] (**Figure 3A**). Performance varied across technology combinations (**Figure 3B**). For Visium–Visium HD, rigid Spateo achieved the highest median PCC, followed by INSPIRE, SPACEL, and STAIR, and also performed strongly across the other gene-expression metrics (**Figure S3**). For Visium–Xenium, rigid Spateo and SLAT with ICP refinement were among the strongest performers, whereas SPACEL, STaCker, and landmark-assisted STalign showed comparatively high median PCC for Visium HD–Xenium. Notably, SPACEL performed competitively on the two Visium HD-containing pairs but showed PCC values concentrated near zero for Visium–Xenium, indicating that performance on one technology combination did not generalize to another. Rankings also varied across gene-expression metrics, emphasizing the importance of considering multiple measures of spatial expression correspondence (**Figure S3**). Computational constraints and incomplete outputs limited method coverage, highlighting the practical challenges of aligning large datasets from high-resolution platforms.

**Figure 3:**
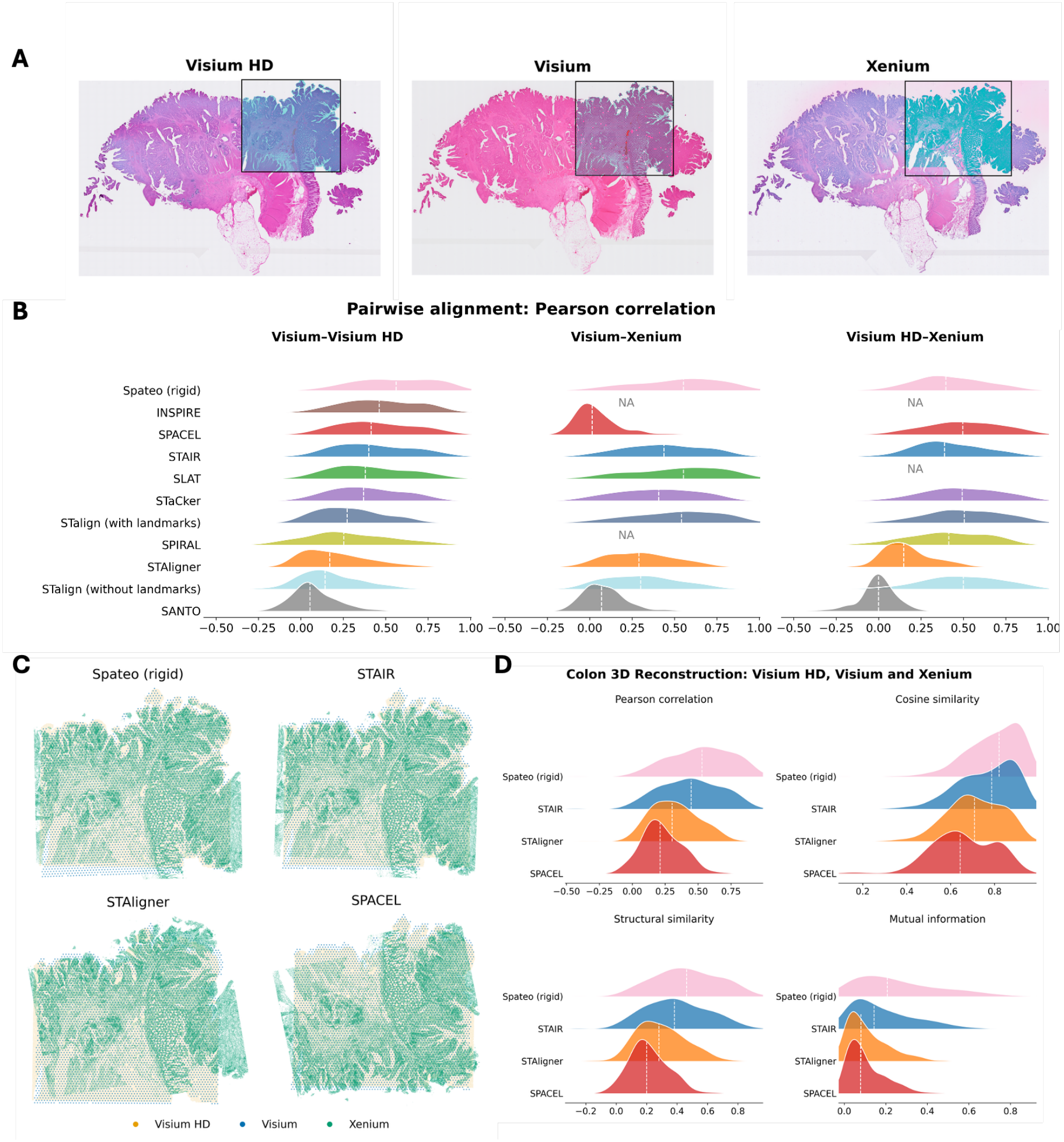
Cross-technology pairwise alignment and three-dimensional reconstruction of human colon cancer tissue profiled with Visium HD, Visium, and Xenium. **(A)** Histological images of the three tissue sections with spatial observation locations overlaid. Black rectangles indicate the spatial extent of the observations used for alignment. **(B)** Distributions of gene-level Pearson correlation coefficients (PCC) following pairwise alignment for Visium–Visium HD, Visium–Xenium, and Visium HD–Xenium. Scores were calculated from grid-aggregated expression profiles, retaining genes with at least 50 nonzero shared grid cells. NA indicates that no eligible scores were available for the corresponding method and technology pair. **(C)** Overlays of the three aligned sections from reconstructions generated by Spateo (rigid), STAIR, STAligner, and SPACEL. **(D)** Distributions of gene-level PCC, cosine similarity, structural similarity index (SSIM), and mutual information (MI) for the three-section reconstructions. In **(B)** and **(D)**, ridgelines show kernel density estimates of the gene-level scores, and white dashed lines indicate medians.

For reconstruction incorporating all three technologies, results were available for rigid Spateo, STAIR, STAligner, and SPACEL (**Figure 3C**,**D**). Rigid Spateo achieved the highest median scores across PCC, cosine similarity, SSIM, and MI, followed by STAIR. Examination of the reconstructed tissue geometry identified an additional limitation of SPACEL: the Xenium section was flipped relative to the other two sections, disrupting correspondence between tissue structures (**Figure 3C**). This contrasted with its competitive VisiumHD–Xenium pairwise performance and demonstrated that strong correspondence for an individual pair did not guarantee a consistent three-section reconstruction.

### 2.4 Simulation-based assessment of alignment

We evaluated alignment methods using a simulated three-dimensional tissue containing six spatial domains, sectioned along three orientations and represented at cell, bin, and spot resolutions (**Figure 4A**,**B**). Sectioning the same simulated sphere along the X, Y, and Z axes generated distinct tissue cross-sections and patterns of domain correspondence, enabling evaluation across different structural contexts within a shared tissue model. These representations provided complementary views of the same underlying tissue architecture. Spatial domains differed in cell-type composition, while individual cell types remained interspersed within domains (**Figure S4**). The unaligned inputs exhibited differences in section position and orientation across the nine resolution–orientation conditions (**Figure S5**).

**Figure 4:**
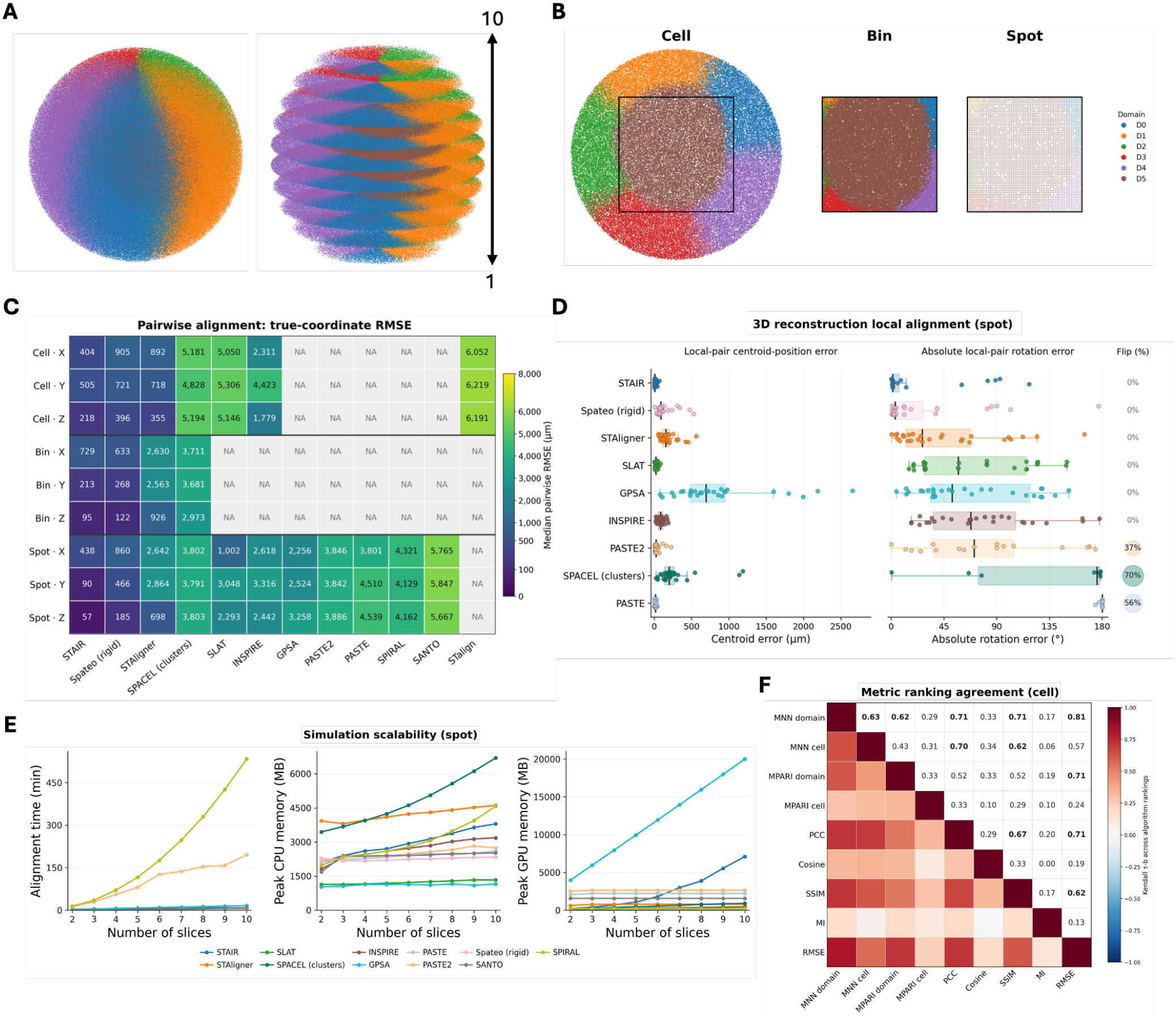
Simulation-based evaluation of spatial alignment and three-dimensional reconstruction. **(A)** Simulated three-dimensional tissue containing six spatial domains and ten sections along the Z orientation. Colors indicate domain identity; spacing between sections is illustrative. **(B)** Cell-, bin-, and spot-level representations of the same simulated section. Black rectangles indicate the capture region for bin- and spot-level observations. **(C)** Heatmap of median pairwise root mean squared error (RMSE), in micrometers, across three resolutions and three sectioning orientations (X, Y, and Z). NA denotes unavailable results. **(D)** Local centroid-position errors, absolute rotation errors, and flip percentages for spot-level 3D reconstructions. Points represent consecutive section pairs pooled across X, Y, and Z. Reflected pairs are excluded from rotation-error distributions. Boxes indicate the interquartile range, center lines indicate medians, and whiskers extend to the most extreme observations within 1.5 times the interquartile range. **(E)** Alignment time, peak CPU memory, and peak GPU memory for spot-level reconstruction with increasing numbers of sections. Points show median values across X, Y, and Z. **(F)** Metric ranking agreement for screened cell-level section pairs, summarized across X, Y, and Z. The heatmap compares domain- and cell-type-based MNN and MPARI, gene-expression-based PCC, cosine similarity, SSIM, and MI, and coordinate-based RMSE.

STAIR and rigid Spateo generally produced the smallest pairwise positional errors relative to the simulated ground truth (**Figure 4C**). STAIR achieved the lowest median RMSE across all cell- and spot-level conditions and two of the three bin-level conditions, while rigid Spateo performed best in the remaining bin-level condition. Their consistently strong relative performance across orientations suggests robustness to the different tissue cross-sections represented in this simulation, although absolute errors varied with orientation and resolution. STAligner was also competitive at cell resolution but showed less consistent performance at coarser resolutions. Other methods exhibited substantially larger errors in several conditions, highlighting that alignment accuracy depended on both the method and the structural context of the input sections. Method coverage decreased at higher observation counts. Only four methods had eligible pairwise results across all nine conditions, whereas a broader set could be evaluated at spot resolution. We restricted this comparison to results obtained without external downsampling, and computational constraints limited method coverage on larger datasets. Spateo’s built-in sparse and chunked computation enabled alignment of the bin-level datasets without external downsampling, while retaining competitive accuracy.

Although RMSE summarizes positional accuracy, it does not distinguish whether an error arises from displacement, rotation, or reflection. We therefore examined local centroid-position errors, residual rotation errors, and relative flips between consecutive sections in the reconstructed 3D stacks (**Figure 4D**). Evaluating each section relative to its predecessor helps distinguish errors at individual section interfaces from deviations inherited from earlier sections. In spot-level reconstructions, STAIR showed small centroid-position errors and rotation errors concentrated near zero, although some consecutive pairs exhibited larger rotations. Rigid Spateo also showed predominantly small errors with several larger deviations. Small centroid errors did not necessarily imply correct orientation: SLAT, INSPIRE, and PASTE included pairs with limited center displacement but substantial rotation errors. This may reflect ambiguity in the correspondences used to estimate alignment, where approximately symmetric tissue geometry can permit substantial rotation around a correctly positioned center. False flips occurred in 70% of consecutive pairs for SPACEL, 56% for PASTE, and 37% for PASTE2. Such reflections preserve within-section distances and can retain similar tissue outlines, potentially allowing incorrect orientations to satisfy alignment objectives when molecular or structural features provide insufficient directional information. The corresponding cell- and bin-level diagnostics showed that these error profiles also varied with resolution (**Figure S6**). The reconstruction RMSE summaries further demonstrated task- and condition-dependent performance (**Figure S7A**). STAIR achieved the lowest local reconstruction RMSE across all spot-level conditions, while STAIR and rigid Spateo alternated as the strongest performers at cell and bin resolutions. However, STAIR showed substantial variation across bin-level conditions, indicating that its strong performance was not uniform across tissue cross-sections. For cross-resolution reconstruction, rigid Spateo consistently achieved the lowest median RMSE across all three orientations, ranging from 113 to 190 micrometers (**Figure S7B**). These results indicate that strong pairwise performance did not translate uniformly across reconstruction tasks and resolutions.

Three-dimensional reconstruction becomes increasingly computationally demanding as both the number of sections and the number of observations per section increase. We assessed scalability by increasing the number of input sections from two to ten and recording alignment time and peak CPU and GPU memory usage (**Figure 4E**; **Figure S8**). We specified four CPU cores and one NVIDIA A100 GPU (80 GB GPU memory), with host-memory allocations of 300 GB across all tested workflows. Runtime generally increased with the number of sections, but the rate of increase differed substantially among methods; peak memory usage did not necessarily increase in parallel with runtime. At spot resolution, rigid Spateo and SLAT had the shortest recorded alignment times, whereas SPIRAL and PASTE2 required approximately 533 and 196 minutes, respectively, at ten sections. At this scale, SPACEL had the highest peak CPU memory usage, approximately 6,700 MB, while GPSA had the highest peak GPU memory usage, approximately 20,010 MB. Fewer methods completed the cell- and bin-level experiments. Spateo’s built-in sparse and chunked computation enabled completion of the bin-level experiments, whereas disabling these options resulted in GPU out-of-memory failures even with two sections. These results highlight the importance of memory-efficient implementations for extending accurate alignment to large, high-resolution tissue reconstructions.

Finally, the simulations allowed us to examine whether alternative evaluation metrics ranked methods consistently with positional accuracy. Domain-based scores depended on the underlying correspondence between sections: even when sections were evaluated at their known simulated positions, MNN accuracy and MPARI could be low when spatial domain correspondence was limited (**Figure S9**). These scores therefore reflect both alignment quality and the availability of corresponding tissue structure, limiting their ability to distinguish registration errors from genuine differences between sections. This distinction is particularly relevant to experimental datasets, where partial overlap and biological variation may occur even under anatomically appropriate alignment. For the metric-agreement analysis, we restricted comparisons to section pairs with sufficient tissue correspondence defined from the simulated ground-truth structure, reducing the influence of intrinsically limited correspondence when comparing algorithm rankings.

Within these screened cell-level pairs, MNN domain accuracy showed the strongest ranking agreement with RMSE, with a summarized Kendall’s tau-b of 0.81 (**Figure 4F**). Domain MPARI and PCC each reached 0.71, followed by SSIM at 0.62 and cell-type MNN at 0.57. Cell-type MPARI, cosine similarity, and MI showed weaker agreement, at 0.24, 0.19, and 0.13, respectively. Agreement also extended across metric categories: MNN domain accuracy agreed with PCC and SSIM at 0.71, while PCC and SSIM agreed at 0.67. Cell-type MNN also showed appreciable agreement with PCC and SSIM, at 0.70 and 0.62, respectively, indicating that these measures captured some common aspects of alignment despite their different relationships with positional accuracy. Across the nine resolution–orientation conditions, MNN domain accuracy and domain MPARI remained positively associated (**Figure S10**). In contrast, the strong relationship between PCC or SSIM and RMSE at cell resolution did not extend to spot resolution. At spot resolution, PCC and SSIM agreed strongly with each other, with tau-b values of 0.80–0.82, but showed weak agreement with RMSE, ranging from 0.07 to 0.16 for PCC and from 0.02 to 0.24 for SSIM. One plausible explanation is that pooling expression from spatially intermixed cell types attenuated the modest compositional differences between simulated domains, reducing the ability of expression-based scores to distinguish positional errors. These findings suggest that metric selection should account for the spatial organization of the data. When cell types are interspersed within compositionally distinct domains, domain-based metrics may be more informative about positional accuracy than cell-type matching, with expression-based metrics providing complementary evidence.

### 2.5 Registration of individual tissue sections to a common coordinate framework

Registering individual tissue sections to a common coordinate framework (CCF) places them within a shared anatomical reference, enabling spatial comparisons across sections and donors. We evaluated STalign [23] and STAIR [22] for registering individual spatial transcriptomic sections to a CCF. Published Allen CCF coordinates for the Zhuang datasets served as the reference for evaluation. STalign registered query sections to the 3D Allen atlas, whereas STAIR used reference sections with known CCF coordinates. Because loading the full reference exceeded the available GPU memory for STAIR, its reference was restricted to a 15-slice window surrounding the expected query location. These experiments therefore evaluated registration with approximate anatomical localization rather than unrestricted localization within the entire brain.

For three held-out sections from Zhuang-ABCA-1, landmark-assisted STalign achieved lower 3D RMSE than STAIR, with errors ranging from 0.168 to 0.434 mm compared with 0.452 to 0.573 mm (**Figure 5A**). Representative reconstructions and coordinate overlays showed that both methods recovered the broad anatomical placement and in-plane organization of the query section (**Figure 5B**). However, axis-specific comparisons revealed residual positional errors despite strong correspondence in the ML and DV coordinates (**Figure 5C**). STAIR assigned an approximately constant AP coordinate to the section, whereas STalign produced spatially varying AP predictions. STalign’s performance depended strongly on initialization and landmark guidance (**Figure S11**). For ABCA-1 slice 66, three-dimensional RMSE increased from approximately 0.297 mm with landmarks and an approximate initial atlas slice to 2.409 mm without landmarks, and to 5.761 mm when landmarks were omitted and the initial atlas slice was randomly selected. These results highlight the importance of anatomical guidance for reliable registration. Such guidance is practical when recognizable landmarks are available, but may be difficult to provide for sections with ambiguous or incomplete anatomical features. Because STAIR uses spatial transcriptomic sections with known coordinates as its reference, we further tested whether this reference that is also registered to the same atlas could be supplied by another donor. Two ABCA-2 query sections were registered using ABCA-1 reference sections (**Figure S12**). We first estimated the approximate AP location of each query section to select a corresponding 15-slice reference window in ABCA-1, and then performed registration using STAIR. Relative to the published CCF coordinates of the ABCA-2 query cells, STAIR achieved 3D RMSE values of approximately 0.236 and 0.374 mm. These results demonstrate the feasibility of transferring spatial localization between donors within a shared coordinate framework, although the use of externally localized reference windows means that this experiment does not establish independent whole-brain localization. Overall, STalign is suited to atlas registration when anatomical landmarks and an approximate initial location are available, whereas STAIR is suited to reference-based localization when spatial transcriptomic sections with known CCF coordinates are available within an approximately identified anatomical region.

**Figure 5:**
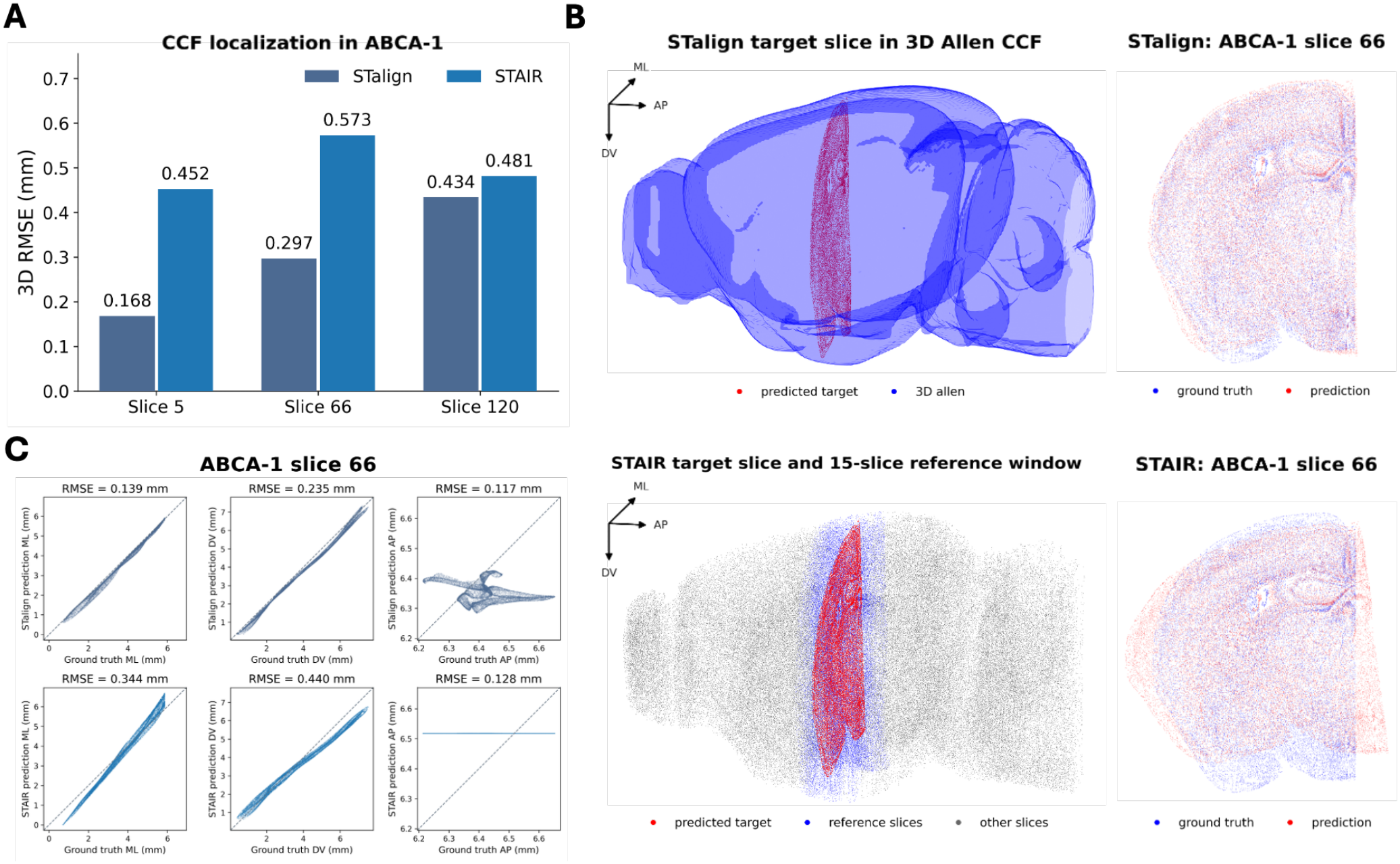
Registration of individual tissue sections to a common coordinate framework. (**A**) Three-dimensional root mean squared error (RMSE) for STalign and STAIR across three query sections from Zhuang-ABCA-1: slices 5, 66, and 120. (**B**) Representative registration of ABCA-1 slice 66 using STalign (top) and STAIR (bottom). Left: STalign-predicted coordinates within the 3D Allen CCF, and STAIR-predicted coordinates within a 15-slice reference window. Right: overlays of predicted and ground-truth cell coordinates in the mediolateral–dorsoventral plane. (**C**) Predicted versus ground-truth coordinates for slice 66 along the mediolateral (ML), dorsoventral (DV), and anteroposterior (AP) axes, shown for STalign and STAIR. Dashed lines indicate perfect agreement (*y* = *x*), and axis-specific RMSE values are reported in millimeters.

## 3 Discussion

We systematically benchmarked 15 spatial alignment and registration algorithms using experimental and simulated spatial transcriptomics datasets. The evaluation covered pairwise alignment, multi-section 3D reconstruction, cross-resolution alignment, and registration to a common coordinate framework. These tasks expose different strengths and limitations and provide a broader basis for method selection. No single method performed best across all settings. Spateo and SPACEL performed strongly across several experimental benchmarks, while STAIR was particularly competitive in simulations with known spatial coordinates (**Figure 6**). However, these advantages depended on the dataset and task, and strong performance in one setting did not consistently generalize to others.

**Figure 6:**
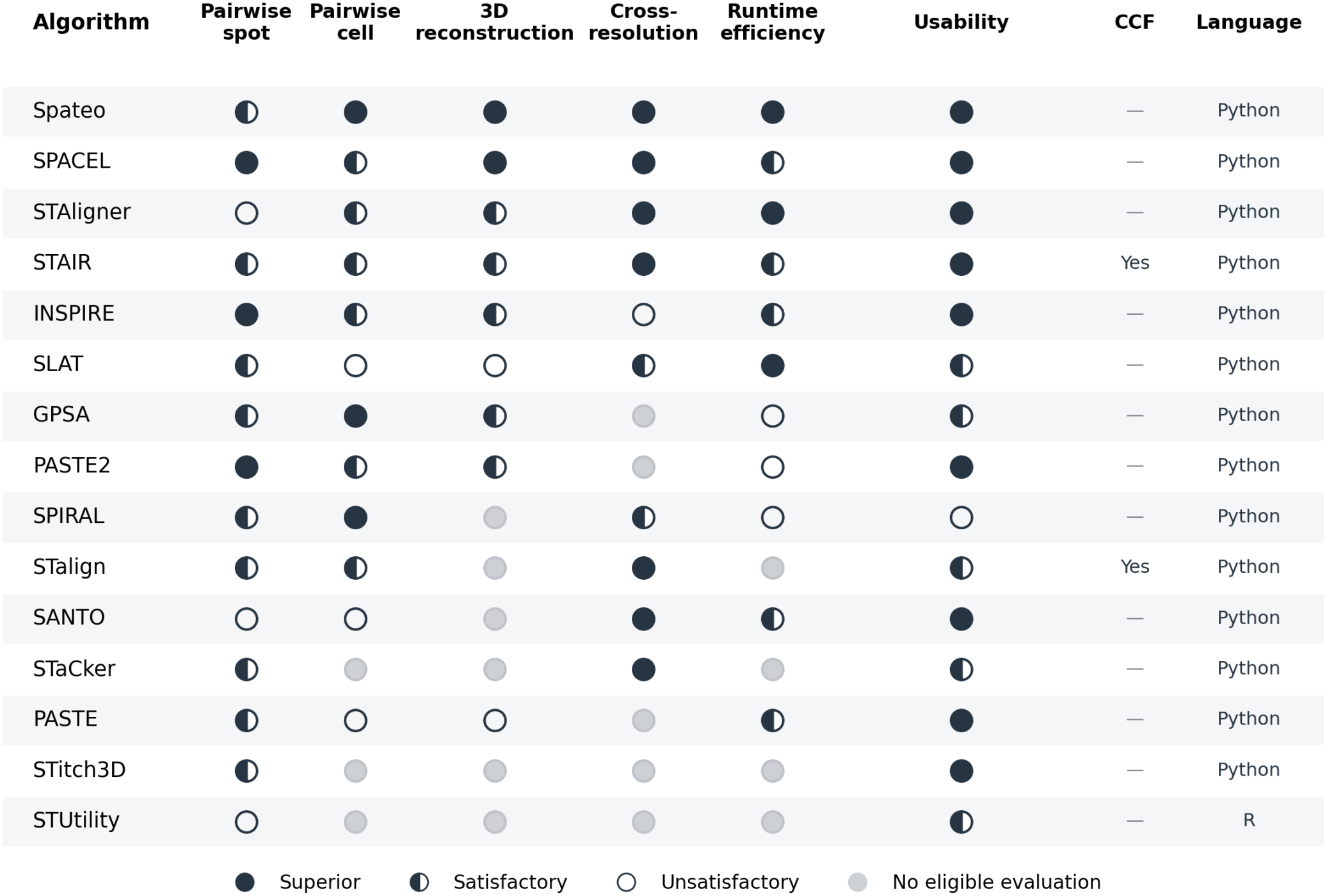
Summary of alignment performance, task coverage, runtime efficiency, and usability across 15 methods. Columns summarize pairwise alignment at spot and cell resolutions, three-dimensional reconstruction, cross-resolution coverage, runtime efficiency, usability, CCF evaluation, and implementation language. Circle symbols indicate the categories shown in the legend, with column-specific rating criteria described in Methods. Gray circles indicate no eligible evaluation, and “Yes” denotes successful evaluation in the CCF task. Spateo denotes its rigid mode, and SPACEL denotes its clustering-based implementation.

A central finding is that alignment performance depends on the biological and technical setting. Several methods performed strongly on human DLPFC, where well-defined cortical layers provide coherent spatial structure, whereas performance was more variable in mouse hypothalamus MERFISH, which combines single-cell resolution, spatially intermixed cell populations, and biological differences between sections. These settings also require different interpretations of evaluation scores: lower cell-type agreement does not necessarily imply poorer geometric alignment, because corresponding tissue regions may preserve cellular composition without matching individual cell identities. Method and metric selection should therefore consider tissue organization, spatial resolution, and the degree of biological correspondence between sections.

Pairwise alignment, multi-section 3D reconstruction, cross-technology registration, and CCF registration are related but distinct computational tasks. Small pairwise errors can accumulate through a reconstructed stack, potentially distorting the overall tissue architecture. Our geometric diagnostics further showed that limited centroid displacement can coexist with substantial rotation or reflection, emphasizing the need to assess orientation as well as positional accuracy. Cross-technology alignment introduces additional differences in observation size, molecular coverage, and tissue sampling, making this task a more stringent test of biological structure recovery. For example, Visium and Visium HD provide transcriptome-wide measurements, whereas Xenium measures a targeted gene panel. Performance varied across these combinations in the human colon benchmark, demonstrating that success on one pairing does not ensure reliable alignment across platforms. CCF registration introduces additional dependencies on anatomical guidance and reference availability. STalign benefited from landmarks and appropriate initialization, whereas the evaluated STAIR workflow used reference sections with known coordinates within an approximately identified anatomical region. Method selection therefore depends on the available reference and the intended registration task.

Evaluation metrics play a critical role in interpreting alignment quality. For example, coordinate-based mutual nearest neighbor accuracy is most appropriate when compared sections are expected to share similar biological structures, such as consecutive serial sections. In settings with true biological variation along the *z*-axis, lower label agreement may reflect genuine changes in tissue composition rather than poor alignment — a distinction that is particularly important for 3D reconstruction tasks. Future benchmarks should develop evaluation strategies that can separate technical misalignment from real biological variation across the z-plane. Simulated datasets with known ground truth are valuable in this regard, though they cannot fully capture the complexity of real tissue sections, including tissue tearing, missing regions, and platform-specific artifacts. Comprehensive benchmarking therefore requires combining simulation-based evaluation with real-world datasets.

Looking forward, future work in 3D spatial alignment could focus on several key directions. First, evaluation metrics should distinguish technical misalignment from biological variation across serial sections while assessing both local correspondence and global tissue structure. Second, algorithms should better accommodate partial overlap, missing tissue regions, nonlinear deformation, and cross-resolution matching. Third, scalable implementations are needed as spatial omics datasets grow in size and resolution. Emerging platforms that profile molecular features directly within three-dimensional tissue volumes may also provide valuable datasets for future benchmarks, enabling assessment of section-to-volume registration and reconstruction against independently measured volumetric organization. Cross-modal approaches incorporating transcriptomics, histology, proteomics, epigenomics, and anatomical atlases could further support more complete spatial tissue models. Finally, standardized benchmark datasets, reproducible workflows, and open-source evaluation toolkits will be essential for fair comparison and continued methodological progress.

Overall, this benchmark provides a systematic evaluation framework for selecting and interpreting tissue alignment methods in spatial omics. Rather than identifying a single universally optimal method, our results suggest that method choice should be guided by the specific biological question, dataset type, tissue structure, spatial resolution, available annotations, and computational constraints. By clarifying the strengths and limitations of current algorithms, this work provides practical guidance for users performing spatial alignment and highlights important opportunities for future method development in 2D and 3D spatial biology.

## 4 Methods

An overview of the algorithms, input requirements, outputs, programming language, software version and source-code availability is provided in Supplementary Table S1. An overview of the datasets, technologies, tissue types, spatial resolutions, annotations and benchmark tasks is provided in Supplementary Table S2.

### 4.1 Overview of tissue alignment and registration algorithms

We reviewed the literature on tissue alignment and registration algorithms available through October 2025 using search terms including spatial transcriptomics, spatial alignment, and registration. Of the 24 surveyed methods, 15 were included in the benchmark. We organize these methods into four computational categories corresponding to Figure 1B: probabilistic spatial warping, optimal transport, morphology-based registration, and graph representation. These categories describe complementary components rather than mutually exclusive algorithm families; methods combining multiple strategies are discussed under the relevant categories. Rigid and nonrigid transformations are distinguished within individual methods.

We used author-released implementations, with method-specific input preparation and computational adaptations where required. Most implementations were run in Python, whereas STUtility was run in R. Software versions, repository links, parameter settings, and workflow adaptations are reported in Supplementary Table S1.

#### 4.1.1 Probabilistic spatial warping

Probabilistic spatial warping treats alignment as an inference problem in which coordinate transformations are estimated from spatial and molecular observations. Depending on the formulation, methods model smooth spatial functions or uncertain correspondences between observations and use these models to estimate transformations into a shared coordinate system. Spatial regularization or prior assumptions constrain the transformations while allowing molecular similarity to inform registration. GPSA [13] uses a two-layer Gaussian-process model: the first layer maps section-specific coordinates into a common coordinate system, and the second models molecular measurements in that shared space. This formulation supports nonlinear alignment of multiple sections and spatial prediction between observed locations. Spateo [19] instead estimates probabilistic correspondences using spatial coordinates and molecular similarity, jointly optimizing coordinate transformations within a probabilistic point-set registration framework. Its alignment module supports rigid and nonrigid registration and provides aligned coordinates for multi-section reconstruction and downstream morphometric analysis.

#### 4.1.2 Optimal transport

Optimal transport approaches represent cross-section correspondence as a transport plan that distributes matching weights between cells or spots. The objective typically balances molecular similarity with preservation of within-section spatial relationships, enabling correspondence estimation without requiring the sections to share an initial coordinate frame. The resulting coupling can guide coordinate transformations, while partial transport formulations allow some tissue regions to remain unmatched. PASTE [6] formulates pairwise alignment as a fused Gromov–Wasserstein optimal transport problem that combines gene-expression dissimilarity with differences in within-section spatial distances. The inferred coupling guides rigid coordinate alignment, enabling serial sections to be assembled into a reconstructed stack. PASTE2 [15] extends this formulation to partial optimal transport, allowing only a specified proportion of tissue mass to be matched and thereby accommodating incomplete overlap or section-specific tissue regions. Optimal transport also forms part of hybrid workflows. SPIRAL [20] uses cluster-aware Gromov– Wasserstein optimal transport to establish correspondences within shared domains and guide coordinate alignment, with cluster assignments supplied by its representation-learning module. STitch3D [25] provides PASTE-based alignment as an alternative to iterative closest point registration during coordinate preprocessing. This establishes the spatial arrangement used by its subsequent three-dimensional graph model, rather than incorporating optimal transport into the graph-learning objective.

#### 4.1.3 Morphology-based registration

Morphology-based registration aligns sections using tissue structure represented by histological images, spatial density maps, or extracted boundaries. Methods estimate transformations by matching image patterns, contours, or geometric landmarks, with molecular information optionally contributing through derived tissue representations. Depending on the transformation model, registration can correct global position and orientation or accommodate local deformation, after which the transformation is applied to the spatial observations. STalign [23] rasterizes spatial observations into density images and estimates affine and diffeomorphic transformations using large-deformation diffeomorphic metric mapping. It supports registration between tissue sections and registration of individual sections to a three-dimensional anatomical atlas. STUtility [26] extracts tissue outlines from histological images and uses iterative closest point registration to align sections, transferring the estimated transformations to spatial observations for joint visualization and three-dimensional tissue assembly. STaCker [21] combines histological images with transcriptomic contour maps derived from expression clusters and uses a neural image-registration model trained with synthetic deformations to map sections into a shared spatial frame. Geometric registration also complements representation-based methods. STAIR [22] refines its embedding-guided initial alignment using tissue and spatial-domain boundary points in an iterative closest point procedure. STitch3D [25] supports iterative closest point registration of section coordinates as an alternative to PASTE-based preprocessing, before constructing its three-dimensional neighborhood graph.

#### 4.1.4 Graph representation

Graph-based approaches represent cells or spots as nodes connected by spatial or molecular relationships and learn features that preserve neighborhood structure across sections. This category also includes broader representation-learning methods that use integrated features or inferred domains to guide geometric registration; alignment of latent representations is distinguished from transformation of physical coordinates. STAligner [24] combines a graph attention autoencoder with triplet learning to identify shared spatial features, with correspondences in the integrated embedding guiding iterative closest point registration. SLAT [17] combines graph convolution with adversarial learning to match spatial observations across datasets, including heterogeneous technologies and modalities. Its cross-section correspondences can subsequently be used to estimate coordinate transformations. SPACEL [18] combines cell-type composition modeling with graph-based learning of shared spatial domains; its Scube module uses domain correspondence and differential evolution to optimize section transformations for three-dimensional reconstruction. INSPIRE [14] combines spatially informed representation learning, non-negative matrix factorization, and adversarial learning to identify shared interpretable factors. Mutual nearest neighbors in its integrated representation provide anchors for rigid registration of adjacent sections. SPIRAL [20] uses graph domain adaptation to separate biological variation from batch-associated variation, producing integrated representations and shared domain assignments for its optimal transport alignment module. SANTO [16] initializes alignment using molecular correspondences and an SVD-based transformation, then learns spatial and molecular neighborhood features using dynamic graph convolutional networks. The resulting correspondence map refines registration for alignment or stitching of partially overlapping datasets. STAIR [22] uses hetero-geneous graph attention with spot- and slice-level mechanisms to learn shared representations and guide initial coordinate alignment; slice-level relationships additionally support relative depth estimation and localization of new sections within an existing spatial transcriptomic reference. STitch3D [25] constructs a three-dimensional neighborhood graph from prealigned sections and learns spatial representations using a graph attention network. With single-cell reference information, this supports identification of three-dimensional spatial domains and estimation of cell-type composition.

#### 4.1.5 Implementation exceptions and reproducibility

All method-specific implementation decisions were recorded before downstream evaluation. For each algorithm, we stored the software version or commit, programming language, script-level entry point, input object format, output files used for evaluation and any deviations from author-recommended or software-default settings. When methods required stochastic initialization, fixed seeds were used where supported by the software. When methods required a reference slice, histology image, spatial graph or shared gene set, these inputs were generated using standardized preprocessing workflows before method execution. These implementation details are summarized in Supplementary Table S1.

### 4.2 Overview of data

For our benchmarking tasks, we use four datasets spanning both spot-based and image-based sequencing technologies. All datasets are normalized and log-transformed when required, depending on the specific algorithm, using Scanpy.

#### 4.2.1 Real world data

##### Human dorsolateral prefrontal cortex (DLPFC) profiled with Visium [29]

This dataset comprises 12 tissue sections from three neurotypical adult donors (two males and one female, aged 30–46 years), sampling the six cortical layers and underlying white matter. Each donor contributed two pairs of spatial replicate sections, with each pair consisting of two directly adjacent, 10 *µ*m-thick serial sections. The second pair was collected 300 *µ*m posterior to the first. We evaluated all six pairwise combinations within each donor, yielding 18 within-donor pairs spanning adjacent and more widely separated sections. Raw Visium data were processed using a standardized quality-control pipeline. Spot-level annotations were incorporated, and only spots with valid annotations located within tissue were retained. Local quality-control outliers were identified using SpotSweeper [30], and spots flagged as outliers in any evaluated category were removed. Genes expressed in fewer than three spots were filtered out, and mitochondrial (MT-) and spike-in (ERCC) features were excluded.

##### Human colorectal cancer dataset profiled with Visium, Visium HD, and Xenium [4]

The human colon dataset consists of spatial transcriptomics data generated using Visium HD, Visium, and Xenium technologies, including both normal adjacent tissue (NAT) and colorectal carcinoma (CRC) samples from five patients. One patient (P2) was profiled using all three technologies, enabling cross-platform comparisons. The Visium HD dataset (16 *µ*m binned outputs) includes five slices from different donors, while the Visium and Xenium datasets each contain three slices from distinct donors. For Visium and Visium HD data, spots were first filtered to retain only those within tissue. Quality control metrics, including total counts, number of detected genes, and mitochondrial gene percentage, were computed using Scanpy. Spatially aware outlier detection was performed using SpotSweeper, identifying local outliers across these metrics. Spots flagged as outliers in any metric were removed. Additionally, genes expressed in fewer than three spots were excluded, and mitochondrial (MT-) and spike-in genes (ERCC, if present) were removed. No additional normalization or log-transformation was applied, as downstream analyses were performed on the filtered count data. For Xenium data, cell-level quality control was performed using median absolute deviation (MAD)-based filtering (4 MADs below the median) on total counts and number of detected genes.

Cells with zero count density or non-zero control probe or codeword counts were excluded. A combined QC flag was used to remove low-quality cells. Spatial coordinates were derived from centroid positions and aligned with histological images. All processed datasets were subsequently used for downstream analysis and cross-technology spatial alignment.

##### Mouse hypothalamic preoptic region [31]

The mouse hypothalamic preoptic region dataset comprises 181 spatially resolved samples derived from 36 animals, each exhibiting distinct behavioral states including naive, parenting, virgin parenting, aggression toward pups or adults, and mating. Gene expression was measured using MERFISH, capturing 135 genes from combinatorial smFISH and 21 additional genes from non-combinatorial rounds. The raw data were first processed in R by replacing missing gene counts with zero and partitioning the dataset into slice-level subsets based on unique Animal ID and Bregma combinations. Cells annotated as Ambiguous were excluded at this stage. For each slice, a Seurat object was constructed using gene expression counts and associated metadata, including spatial coordinates, cell class, and neuron cluster identity. These objects were then converted to SingleCellExperiment and sub-sequently to SpatialExperiment format, and exported as individual h5ad files. In the subsequent Python preprocessing step, control probes were removed, and quality control metrics, including total transcript counts and number of detected genes per cell, were computed. Cells with zero total counts or zero detected genes were flagged as low quality. As the dataset was already normalized, only a log-transformation was applied. Mouse hypothalamic preoptic region. The mouse hypothalamic preoptic region dataset consists of 181 individual samples from 36 mice. Probe genes are removed, and cells with a total gene count or number of detected genes equal to 0 are excluded.

##### Mouse Primary Motor Cortex [28]

The dataset was already preprocessed, normalized and log-transformed. The preprocessed `.h5ad` file includes gene expression data along with spatial coordinates and associated metadata, such as slice identifiers and spatial domain annotations. No additional normalization or filtering was applied prior to downstream analysis and visualization.

##### Mouse whole-brain MERFISH data for CCF registration [32]

We used the Zhuang-ABCA-1 and Zhuang-ABCA-2 datasets from the Allen Brain Cell Atlas, representing two adult mouse donors profiled by MERFISH. The released datasets contain 147 and 66 coronal sections, respectively, sampled at nominal intervals of 100 and 200 *µ*m, with expression measurements for 1,122 genes. Each dataset includes native spatial coordinates, cell-type annotations, and published coordinates registered to the Allen Mouse Brain Common Coordinate Framework version 3 (CCFv3). For STalign, we additionally obtained the Allen CCF average anatomical template at 25 *µ*m isotropic resolution from the Allen-CCF-2020 release as the three-dimensional registration reference. STAIR instead used spatial transcriptomic reference sections with known CCF coordinates. Five query sections were selected: ABCA-1 slices 5, 66, and 120 (Zhuang-ABCA-1.014, .076, and .132) and ABCA-2 slices 21 and 39 (Zhuang-ABCA-2.025 and .046). These sections supported evaluation of within-donor registration, cross-donor reference mapping, and initialization sensitivity. Published CCF coordinates were treated as registration-derived reference positions rather than independently measured anatomical ground truth.

#### 4.2.2 Synthetic data

Synthetic benchmark datasets were generated using ALBIS with its default settings. The simulated tissue was spherical, with a diameter of 12 mm, and contained 600,000 cells, six spatial domains (D0–D5), eight cell types (type1–type8), and 556 genes. The domain structure comprised a central region surrounded by five peripheral regions with irregular boundaries. Domains differed in cell-type composition, but cell types were spatially interspersed within each domain rather than forming separate, sharply defined regions.

The same underlying simulated RNA molecules were represented at three spatial resolutions: individual cells, square bins with an 8-micrometer side length, and spots with a 55-micrometer diameter and 100-micrometer center-to-center spacing. Bin- and spot-level observations covered a central 6.5 mm by 6.5 mm capture window, whereas cell-level observations covered the full tissue. The tissue was sectioned along the X, Y, and Z orientations, with ten sections per orientation, yielding nine resolution–orientation datasets. Each orientation contained 600,000 cell-level, 6,609,690 bin-level, and 43,560 spot-level observations across its ten sections, before algorithm-specific filtering.

Each dataset included gene-expression counts, section identifiers, spatial-domain and cell-type annotations, and ground-truth spatial coordinates. Unaligned coordinates with section-specific rotations and translations were provided as alignment inputs, and section-specific expression variation was included. These datasets supported pairwise alignment, within-resolution 3D reconstruction, and cross-resolution reconstruction benchmarks. Each ten-section dataset provided 45 distinct section pairs, including both consecutive and non-consecutive pairs.

### 4.3 Evaluation metrics

#### 4.3.1 Cell- and domain-based accuracy metrics

To evaluate whether aligned slices preserve local cellular and tissue-domain relationships, we computed a mutual nearest neighbor (MNN)-based accuracy metric in the aligned coordinate space. For each pair of aligned slices, spots or cells were represented by their aligned spatial coordinates and MNN pairs were identified between slices using approximate nearest-neighbor search. To ensure sufficient correspondence coverage, the number of nearest neighbors *k* was iteratively increased until at least 80% of observations in the smaller slice participated in at least one MNN match, or a predefined maximum value of *k* was reached. Using manually annotated cell-type or tissue-domain labels, accuracy was defined as the proportion of MNN pairs for which the labels of the matched observations agreed across slices,

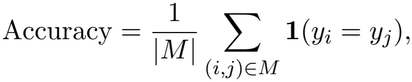

where *M* denotes the set of MNN pairs and *y*_*i*_ and *y*_*j*_ are the labels of the matched observations. Coverage was defined as

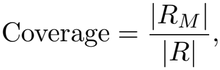

where *R* denotes the observations in the smaller slice and *R*_*M*_ denotes the subset participating in at least one MNN pair. Higher accuracy indicates better preservation of local biological structure following alignment, while higher coverage indicates a larger fraction of the tissue that can be reliably matched between slices.

#### 4.3.2 Gene-based evaluation metrics

When aligning consecutive tissue slices from biological samples, we assume that gene expression is spatially conserved across slices: under a correct alignment, cells with similar transcriptomic profiles should occupy spatially proximate locations. Gene expression-based similarity metrics are particularly well-suited for cross-technology benchmarking — for example, comparing Visium against Xenium — because cluster-based metrics such as ARI and MPARI require a shared cell type label space, which is difficult to establish when comparing technologies with fundamentally different gene panels and resolutions. The calculation of gene similarity metric is based on previous work by [11] [18].

To compute these metrics, we first overlay a regular square grid on the intersection of the two slices’ coordinate bounding boxes, using the aligned coordinate system shared across slices. Grid cell edge length is technology-pair-specific to account for differences in spatial resolution across acquisition platforms. The same grid is applied to both slices simultaneously: spots, bins, or cells from each slice are independently assigned to the grid cell they fall within, and spots outside the intersection bounding box are excluded. A grid cell is retained only if both slices meet a minimum occupancy threshold of at least 50% of the smaller of the two slices’ mean per-cell occupancy, ensuring that retained cells contain sufficient observations from both technologies.

For each retained grid cell, we compute the mean expression of each gene across all spots, bins, or cells assigned to that grid cell, independently per slice. This produces two vectors of length `n_grids` per gene, one per slice, representing the spatial expression profile at matched physical locations. Metrics are computed in two variants. The first uses all retained grid cells. The second restricts computation to the subset of grid cells where mean expression is non-zero in both slices simultaneously, i.e. grid cells *ω* satisfying *x*^(*ω*)^ *>* 0 and *x*^(*ω*)^ *>* 0, thereby reducing the influence of zero-inflation on the similarity estimates. Although metrics were computed for all genes, downstream visualizations and summaries of the non-zero grid-cell variant included only gene-level metric estimates where at least 50 grid cells showed non-zero mean expression in both slices (unless otherwise specified). Genes included in the analysis are restricted to the intersection of gene panels across the compared slices. We compute four metrics per gene: Pearson correlation coefficient (PCC), cosine similarity, structural similarity index (SSIM), and mutual information (MI). Genes with zero variance are assigned `NaN` for PCC and MI; genes with zero vector norm are assigned `NaN` for cosine similarity; and genes with zero maximum expression are assigned `NaN` for SSIM. Higher values across these metrics indicate greater concordance of spatial expression profiles at matched locations, which under the spatial conservation assumption reflects better alignment quality.

For multi-slice datasets, non-zero gene similarity metrics were computed independently for each pair of consecutive slices and averaged across slice pairs. Pairwise estimates based on fewer than 50 overlapping grid cells with non-zero mean expression in both slices were excluded from the averaging procedure.

For SLAT, we compute two versions of the gene similarity metrics. The first (ICP based SLAT) follows the standard grid-based evaluation after applying the final SLAT alignment, which includes the ICP refinement step implemented within SLAT. The second uses the cell-to-cell correspondences inferred by SLAT directly. In this matched-cell evaluation, we extract the expression profiles of each matched cell pair and construct paired gene-wise expression vectors across all SLAT matches. Gene similarity metrics are then computed on these matched vectors rather than on grid-aggregated expression profiles.

#### 4.3.3 Cosine similarity

Cosine similarity measures the directional concordance between two spatial gene expression profiles, independent of their absolute magnitudes. For a given gene, let **x**_*i*_ and **x**_*j*_ denote the vectors of grid-aggregated mean expression across *L* retained grid cells in slices *i* and *j*, respectively. Cosine similarity is defined as

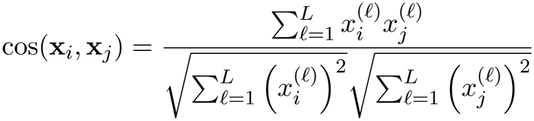

Values close to 1 indicate that the gene has a similar spatial expression pattern across the two aligned slices, even if the overall expression magnitude differs between slices. Values close to 0 indicate weak directional agreement between the two spatial profiles. Cosine similarity is therefore useful for cross-technology comparisons, where absolute expression levels may differ due to platform-specific sensitivity, capture efficiency, or resolution.

#### 4.3.4 Mutual information

Mutual information (MI) quantifies the degree of statistical dependence between two variables. For two random variables *X* and *Y* with joint probability distribution *p*(*x, y*) and marginal distributions *p*(*x*) and *p*(*y*), MI is defined as

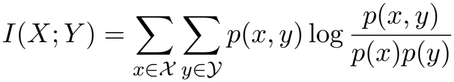

In our setting, *X* and *Y* represent the grid-aggregated mean expression of a given gene across retained grid cells in slices *i* and *j*, respectively. Higher MI values indicate stronger statistical dependence between the two spatial expression profiles at matched locations. MI is estimated using a *k*-nearest-neighbour regression approach via `scikit-learn`’s `mutual_info_regression`.

#### 4.3.5 Pearson correlation coefficient

Pearson correlation coefficient (PCC) measures the linear concordance between two spatial gene expression profiles. For a given gene, let **x**_*i*_ and **x**_*j*_ denote the vectors of grid-aggregated mean expression across *L* retained grid cells in slices *i* and *j*, respectively. PCC is defined as

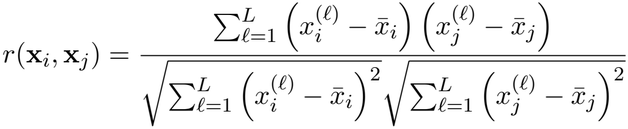

Here, 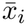 and 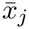 denote the mean expression values across retained grid cells for slices *i* and *j*, respectively. Values range from −1 to 1, where 1 indicates perfect positive linear concordance, 0 indicates no linear association, and −1 indicates perfect negative linear concordance. PCC is computed using `scipy.stats.pearsonr`.

#### 4.3.6 Structural similarity index

Structural similarity index (SSIM) measures concordance between two spatial gene expression profiles by jointly comparing their mean expression, variance, and covariance structure. For a given gene, let **x**_*i*_ and **x**_*j*_ denote the grid-aggregated mean expression vectors across retained grid cells in slices *i* and *j*, respectively. SSIM is defined as

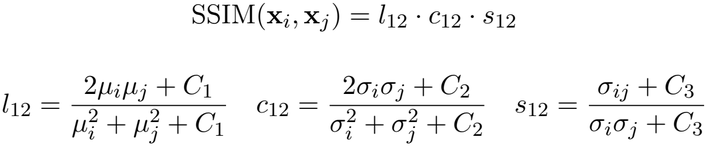

Here, *µ*_*i*_ and *µ*_*j*_ denote the mean expression values, *ε*_*i*_ and *ε*_*j*_ denote the standard deviations, *ε*_*ij*_ denotes the covariance, and *C*_1_, *C*_2_, and *C*_3_ are stabilisation constants with *C*_3_ = *C*_2_*/*2. Prior to computation, each vector is normalised by its maximum value, constraining values to [0, 1]. Higher SSIM values indicate greater structural concordance of the gene’s spatial expression profile across the two aligned slices.

#### 4.3.7 Mean Pairwise Adjusted Rand Index (MPARI)

To quantify the quality of the 3D reconstructions, we report the **Mean Pairwise Adjusted Rand Index** (MPARI), a label-transfer metric that measures how well an alignment preserves cell-type identity across slices. Let *S* = *{S*_1_,. .., *S*_*N*_ *}* denote the *N* aligned slices, each carrying post-alignment spatial coordinates 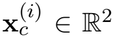 and ground-truth spatial-domain labels 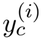 for every cell *c* ∈ *S* . For an ordered pair (*i, j*) we transfer labels from a source slice *i* to a target slice *j* by majority vote over the *k* nearest source neighbours in the aligned coordinate space:

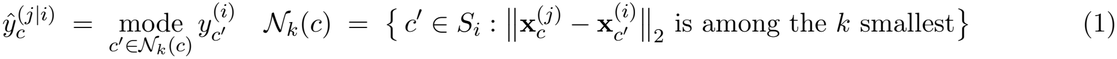

with ties in the majority vote broken in favour of the closer neighbour. Agreement between the transferred labels **ŷ** ^(*j*|*i*)^ and the reference labels **y**^(*j*)^ is quantified by the chance-corrected Adjusted Rand Index:

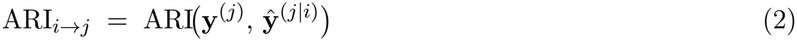

To remove direction asymmetry, we average the two directions of every unordered pair,

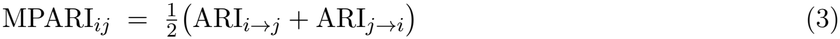

and report the mean over all 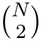 unordered pairs:

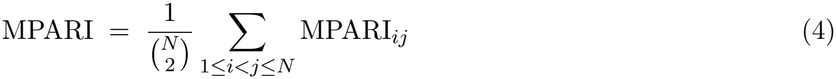

Using ARI rather than raw label-transfer accuracy makes the metric **(i)** invariant to arbitrary permutations of cluster IDs across methods, so that methods whose domain identities are clustered independently are still comparable; **(ii)** robust to severe class imbalance, since each pair of cells contributes independently to the index; and **(iii)** centred at zero under chance assignment, so that an unaligned reconstruction cannot accrue a positive score from base-rate matches alone. Aggregating over *all* slice pairs, rather than only consecutive ones, additionally rewards alignments that remain globally consistent across the entire stack and penalises methods whose registration quality decays with anatomical distance.

In all experiments we used *k* = 5, Euclidean distance, and the exact kNN implementation in `scikit-learn`. To keep the 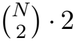 kNN fits tractable, each slice was uniformly subsampled to at most 8,000 cells (random seed 42) prior to neighbour search. Per-method MPARI distributions are reported as the collection of pair scores *{*MPARI_*ij*_*}*_*i<j*_ and summarised by their median.

#### 4.3.8 Relative mean squared error

For simulated datasets, relative alignment accuracy was evaluated using an anchor-corrected mean squared error (MSE). A similarity transformation was estimated from the reference slice:

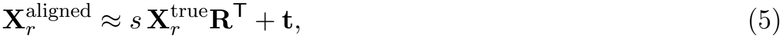

where *s*, **R**, and **t** denote the scaling factor, rotation matrix, and translation vector, respectively. The transformation was then applied to the ground-truth coordinates of the moving slice:

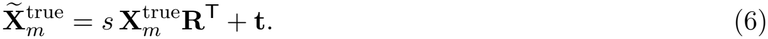

The relative MSE was calculated as

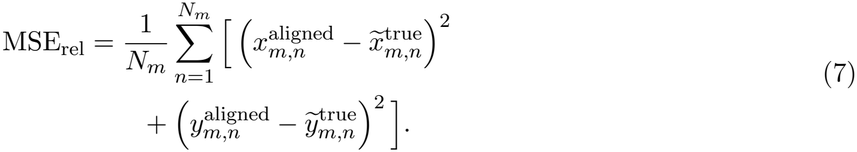

Lower relative MSE indicates more accurate recovery of the moving slice relative to the reference after accounting for global rotation, translation, and isotropic scaling.

#### 4.3.9. Reference normalization for local geometric evaluation

We evaluated local geometric accuracy between consecutive sections in the simulated three-dimensional reconstructions. Aligned observations were matched to their ground-truth coordinates by observation identifiers, using the retained-observation set shared with the RMSE evaluation. Ground-truth coordinates were projected onto the appropriate section plane: (*y, z*), (*x, z*), and (*x, y*) for sections cut along the *x, y*, and *z* axes, respectively.

For each consecutive pair (*i, i*+1), section *i* served as the reference. Let *X*_*i*_ and *Y*_*i*_ denote its matched ground-truth and reconstructed coordinates, respectively, with observations represented as rows. We estimated a similarity transformation by minimizing

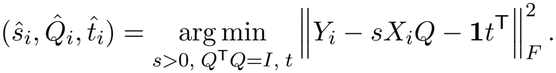

Here, *s* is an isotropic scale factor, *Q* is an orthogonal matrix, and *t* is a translation vector. Reflections were permitted in this reference transformation to remove the arbitrary handedness of the shared coordinate frame. Its inverse was applied to both sections; the corrected moving-section coordinates were

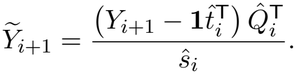

All local errors were then evaluated against *X*_*i*+1_. Thus, each pair was assessed independently of deviations inherited from earlier sections. The reference fitting residual was recorded to identify departures from a similarity transformation, particularly for nonrigid outputs.

#### 4.3.10 Local centroid-position error

Centroid-position error measures displacement of the moving section’s center after reference normalization. For *n* matched moving-section observations, it was calculated as

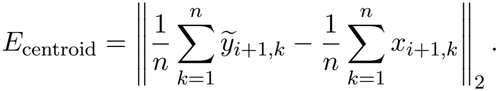

Errors were reported in micrometers. This quantity measures center displacement rather than pointwise positional accuracy and can therefore remain small despite substantial rotation or reflection.

#### 4.3.10 Local rotation error

We estimated the residual orientation of each corrected moving section by fitting a similarity transformation from *X*_*i*+1_ to 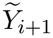 using the same least-squares formulation. Let *R* denote the resulting orthogonal matrix. For orientation-preserving fits, where det(*R*) *>* 0, the signed residual angle under the row-coordinate convention was

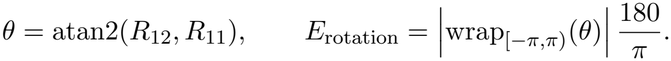

Absolute rotation errors were reported in degrees, ranging from 0 to 180. Pairs with relative reflections were excluded from the rotation-error distribution and evaluated separately. For nonrigid outputs, this angle describes the best-fitting overall orientation rather than local deformation.

#### 4.3.12 Relative flip frequency

A relative flip was identified when the best-fitting orthogonal transformation between the moving section’s ground-truth coordinates and its reference-normalized reconstruction reversed orientation:

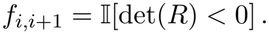

Because the reference section’s transformation was removed first, this detects a difference in handedness between consecutive sections rather than a reflection shared by the entire reconstruction. Correspondences were established by observation identifiers rather than tissue-outline similarity.

For each method and resolution, the flip percentage was

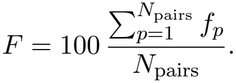

Complete coverage across the three orientations contributed 27 consecutive pairs, comprising nine pairs per orientation. Reflected pairs remained included in centroid-position and RMSE evaluations.

#### 4.3.13 Ground-truth spatial tissue correspondence

We quantified intrinsic tissue correspondence independently of algorithm outputs using simulated ground-truth coordinates and domain labels. For each section pair, coordinates were projected onto the section plane and assigned to a shared square grid with a side length of 300 *µ*m. Let *G*_*ij*_ denote the grid cells occupied by both sections, *n*_*ig*_ the number of observations from section *i* in grid cell *g*, and *p*_*igc*_ the proportion assigned to domain *c*.

The weighted probability of matching domain labels within shared grid cells was

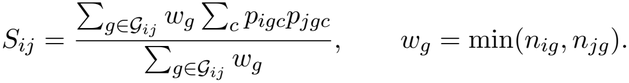

This represents the probability that independently sampled observations from the two sections share a domain label within a corresponding grid cell. Weighting by the smaller observation count limits the contribution of grid cells with strongly unequal sampling densities.

To account for agreement attributable to overall domain composition, we calculated

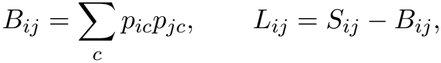

where *p*_*ic*_ is the proportion of domain *c* across all retained observations in section *i*. The spatial correspondence lift *L*_*ij*_ measures excess local domain agreement above this global-composition baseline. Positive values indicate spatial correspondence beyond that expected from domain proportions alone; the score is not a percentage of correctly aligned observations. Analogous quantities were calculated using cell-type labels.

#### 4.3.14 Pair screening for metric-agreement analysis

Pairs were screened using ground-truth tissue properties without inspecting algorithm outputs or evaluation scores. In addition to *S*_*ij*_ and *L*_*ij*_, we calculated projected spatial overlap, global domain-composition overlap, and effective domain diversity:

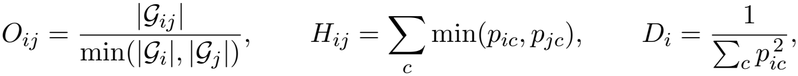

where *G*_*i*_ denotes all occupied grid cells in section *i*. A pair was retained when

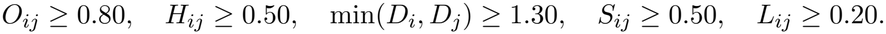

These thresholds selected pairs with sufficient projected overlap, shared domain content, domain diversity, and spatial correspondence for interpretable metric comparisons. For comparisons across resolutions, we retained orientation–pair combinations satisfying all criteria at cell, bin, and spot resolutions. This yielded 45 combinations: 10 for the *x* orientation, 10 for *y*, and 25 for *z*. These screening criteria were applied to the metric-agreement analysis, rather than to the all-pair RMSE benchmark or consecutive-pair reconstruction diagnostics.

#### 4.3.15 Compute benchmarking

All scalability experiments were run under a fixed compute allocation to ensure comparable runtime and memory measurements across methods. Each run was submitted to the GPU partition with 4 CPU cores, a maximum of 300 GB RAM, and one NVIDIA A100 GPU with 80 GB memory. Runtime and memory usage were measured only for the alignment-required steps, excluding data loading, preprocessing, plotting, and other downstream operations. Wall-clock time and peak CPU resident memory were measured using a custom benchmarking implementation based on `psutil`. Peak GPU memory was measured using Py-Torch CUDA memory profiling utilities by resetting peak memory statistics before each run and recording `torch.cuda.max_memory_allocated()`.

#### 4.3.16 Summary ratings across benchmarking tasks

Figure 6 summarizes task-specific performance, cross-resolution coverage, runtime efficiency, and usability using superior, satisfactory, and unsatisfactory categories, represented by filled, half-filled, and open circles, respectively. Gray circles indicate that no eligible evaluation was available; they do not necessarily imply that a method cannot perform the task.

For pairwise spot-level alignment, methods were ranked by median cortical-layer MNN agreement across DLPFC section pairs. Pairwise cell-level alignment used median cell-type MNN agreement across mouse hypothalamus MERFISH pairs, and three-dimensional reconstruction used median spatial-domain MPARI across consecutive mouse motor cortex sections. For each column, higher scores indicated better performance. Among *N* eligible methods, the superior and unsatisfactory categories corresponded approximately to the highest- and lowest-performing 20%, respectively, with the remaining methods classified as satisfactory. Specifically, each tail contained *k* = min(⌈ 0.2*N ⌉*, ⌊ *N/*2 ⌋) ranking positions. Ties received average ranks and were not separated across categories.

Cross-resolution ratings represented coverage rather than alignment accuracy. Methods with eligible gene-expression evaluation results for all three technology combinations—Visium–Visium HD, Visium–Xenium, and Visium HD–Xenium—were classified as superior; those with results for two or one combination were classified as satisfactory or unsatisfactory, respectively. Methods without eligible results for any combination were shown in gray.

Runtime efficiency was evaluated using simulated spot-level reconstruction workloads containing two to ten sections. For each method and section count, runtime was summarized by the median across the three orientations. Methods with complete coverage were ranked at each section count, with shorter runtimes receiving better ranks, and these ranks were averaged across section counts. The resulting ordering was categorized using the same approximate 20%–60%–20% rule.

Usability ratings reflected our experience installing and applying each method. Superior corresponded to easy: a publicly available implementation with clear installation instructions and sufficient documentation to run the relevant workflow. Satisfactory corresponded to medium: an implementation requiring additional dependency configuration, input preparation, or clarification of incompletely documented steps. Unsatisfactory corresponded to hard: substantial troubleshooting or code modification was required, or no implementation was readily available.

CCF evaluation was recorded separately as “Yes” for methods successfully evaluated in the specified localization task; a dash indicated no such evaluation and was not interpreted as evidence of technical in-capability. The language column reported the implementation language. Spateo referred to its rigid mode, and SPACEL to its clustering-based implementation. Methods were ordered by decreasing numbers of superior, then satisfactory, then unsatisfactory ratings across the three accuracy columns, cross-resolution coverage, and runtime efficiency, excluding usability; remaining ties were resolved alphabetically.

## 4.4 Data availability

The raw spatial transcriptomic data used in this study is available from the following studies:

1. Mouse hypothalamic preoptic region dataset, sequenced with Merfish technology: https://datadryad.org/dataset/doi:10.5061/dryad.8t8s248
2. 12 slices of human DLPFC 10X Visium data set: https://bioconductor.org/packages/release/data/experiment/html/spatialLIBD.html
3. 33 slices of the mouse primary motor cortex sample: https://zenodo.org/records/8316334
4. Human colorectal cancer sequenced with Visium, Visium HD and Xenium: https://www.10xgenomics.com/platforms/visium/product-family/dataset-human-crc
5. Mouse whole-brain MERFISH datasets (Zhuang-ABCA-1 and Zhuang-ABCA-2), including registered CCF coordinates, from the Allen Brain Cell Atlas: https://alleninstitute.github.io/abc_atlas_access/notebooks/zhuang_merfish_tutorial.html

### 4.5 Code availability

Code to reproduce all preprocessing, analyses, and figures in this manuscript is available on GitHub at https://github.com/JianingYao/alignment_benchmark_paper.

## 4.6 Acknowledgments, Funding, Authorship Contributions

**Acknowledgments**

We acknowledge the Joint High Performance Computing Exchange (JHPCE) at the Johns Hopkins Bloomberg School of Public Health for computational support, and members of the Hicks lab for their valuable feedback. We thank Haowen Zhou for his help with the mouse hypothalamus dataset. Furthermore we acknowledge the use of Artificial Intelligence to refine the language of the manuscript and assist with coding. The authors take full responsibility for its content and integrity.

## Author Contributions Statement

- Conceptualization of benchmark study: JY, SCH
- Investigation and review of existing papers: JY, JM, GG, VP, AN, SL
- Data aggregation and preprocessing: JY, JM, GG, VP, AN
- Formal analysis: JY, JM, GG, VP, AN
- Writing – original draft: JY, JM, GG, VP, AN, SCH
- Writing – review & editing: all co-authors
- Visualization: JY, JM, GG, VP, AN
- Supervision: SCH
- Funding acquisition: SCH, KRM, KM, DG, LY
- Project administration: SCH, KRM, KM, DG, LY

