## Supplementary materials for "A systematic benchmark evaluation of algorithms for 3D tissue alignment and registration with spatial omics"

#### Contents

1. **Supplemental Tables S1-S2**
2. **Supplemental Figures S1-S12**

### Supplementary Tables

**Supplementary Table S1: Summary of all 3D tissue alignment and registration methods used in each evaluation of this benchmark**. The table includes the name of the method, input, output, pre-processing steps for each method that we applied, the programming language, assumptions about the method, the download date, software version number, and link to software package.

**Supplementary Table S2: Summary of all datasets used in each evaluation of this benchmark**. The table includes the names, protocols, source (GEO accession numbers or links to download) and cell details of each dataset.

### Supplementary Figures

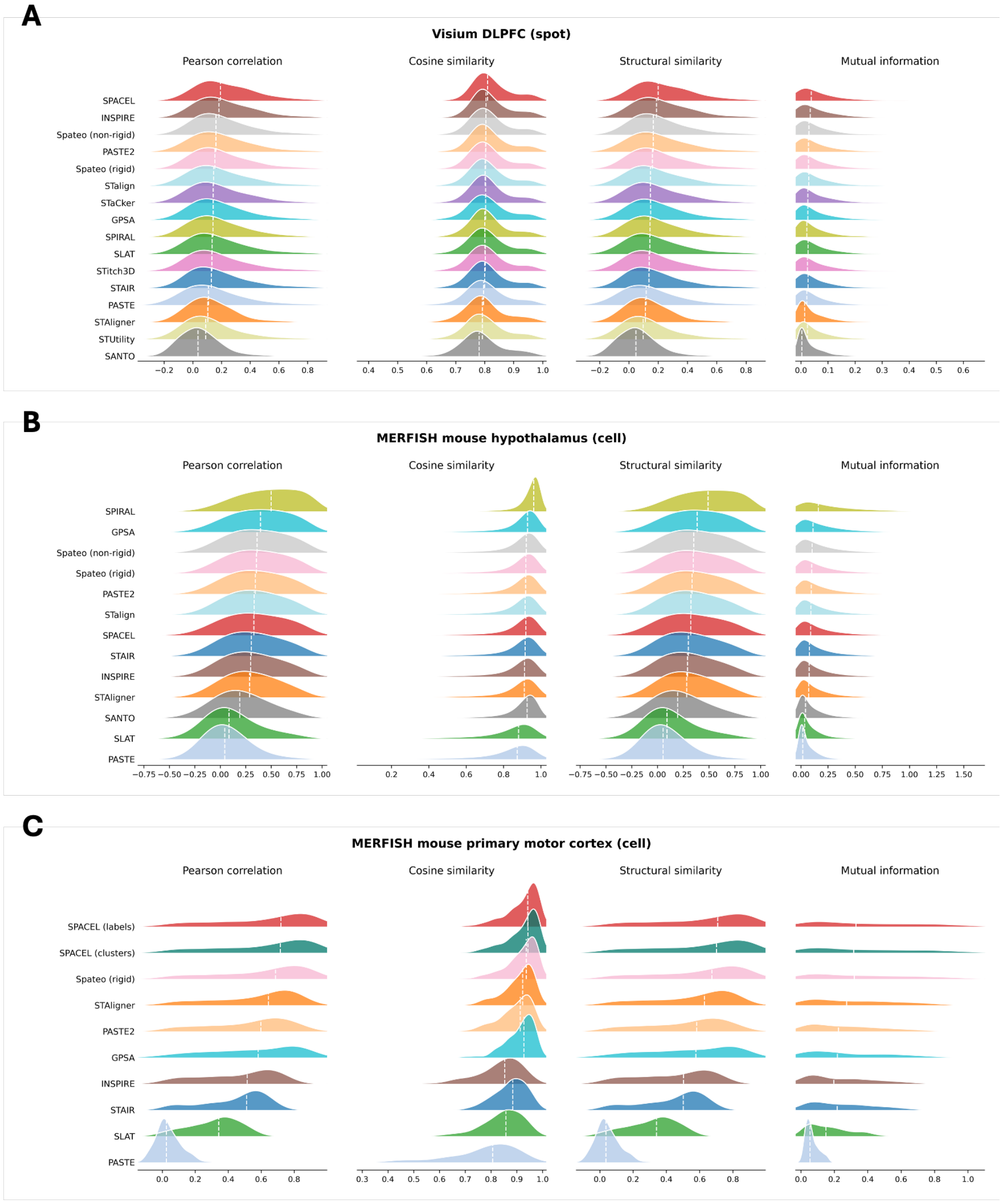


**Supplementary Figure S1: Gene-expression similarity across alignment methods for pairwise alignment and three-dimensional reconstruction.** **(A)** Pairwise alignment of human DLPFC Visium sections. **(B)** Pairwise alignment of mouse hypothalamus MERFISH sections. **(C)** Three-dimensional reconstruction of mouse primary motor cortex MERFISH sections. Columns show distributions of gene-level Pearson correlation coefficient (PCC), cosine similarity, structural similarity index (SSIM), and mutual information (MI).

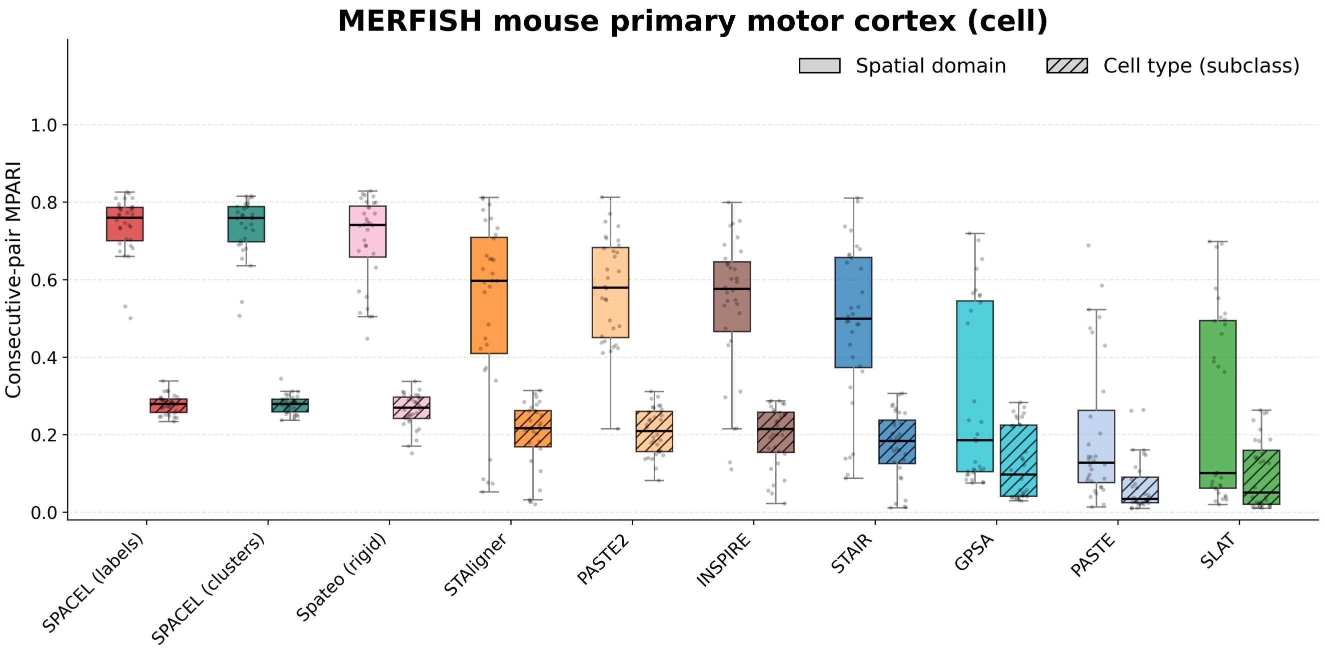


**Supplementary Figure S2: Spatial-domain and cell-type MPARI for three-dimensional reconstructions of mouse primary motor cortex MERFISH data.** Paired boxplots show MPARI distributions across 32 consecutive section pairs for each alignment method. Solid boxes represent spatial-domain annotations, and hatched boxes represent cell-type subclass annotations. Points indicate individual section pairs. Boxes indicate the interquartile range, center lines indicate medians, and whiskers extend to the most extreme observations within 1.5 times the interquartile range.

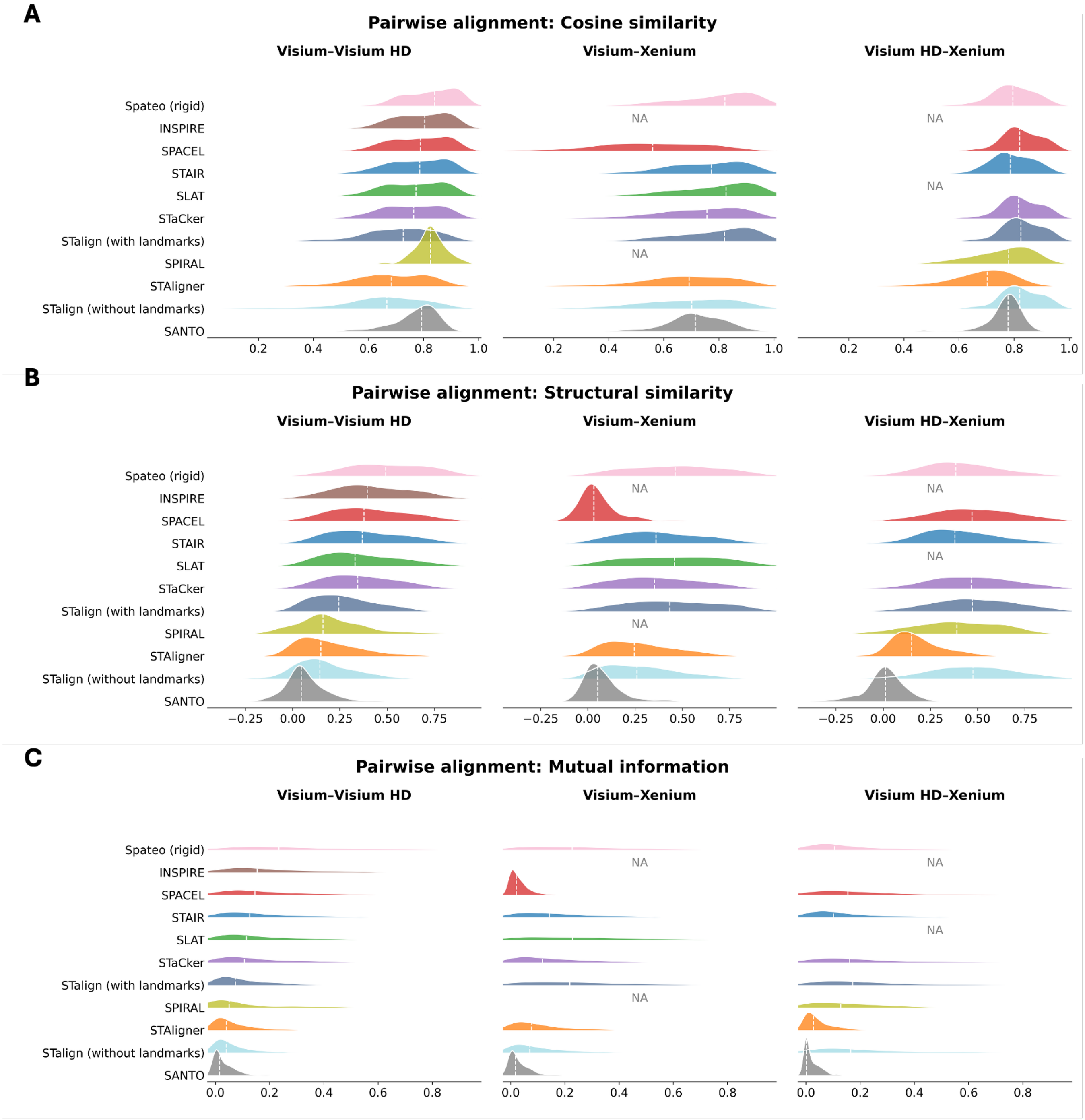


**Supplementary Figure S3: Gene-expression similarity following cross-technology pairwise alignment of human colon cancer spatial transcriptomics datasets.** Distributions of gene-level **(A)** cosine similarity, **(B)** structural similarity index (SSIM), and **(C)** mutual information (MI) are shown across alignment methods. Columns correspond to Visium–Visium HD, Visium–Xenium, and Visium HD–Xenium, respectively. Colored ridgelines represent score distributions, and white dashed lines indicate medians. NA denotes unavailable eligible scores.

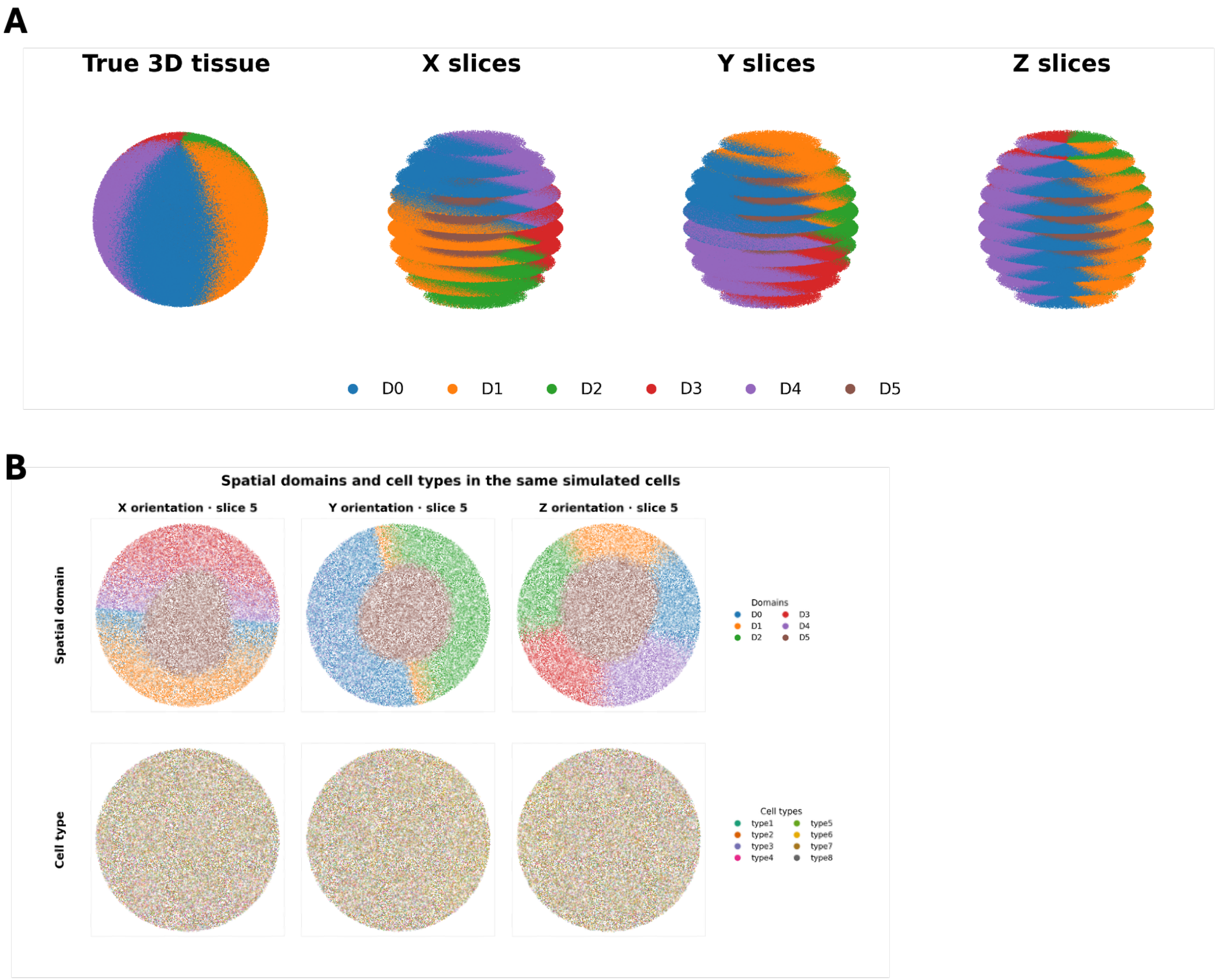


**Supplementary Figure S4: Spatial domains, sectioning orientations, and cell-type distributions in the simulated tissue.** **(A)** Ground-truth three-dimensional tissue and ten sections along each of the X, Y, and Z orientations, colored by spatial domain. Spacing between sections is illustrative. **(B)** Representative cell-level sections with slice ID 5 from each orientation. The same cells at identical coordinates are colored by spatial domain in the top row and by cell type in the bottom row. Columns correspond to X, Y, and Z orientations, respectively.

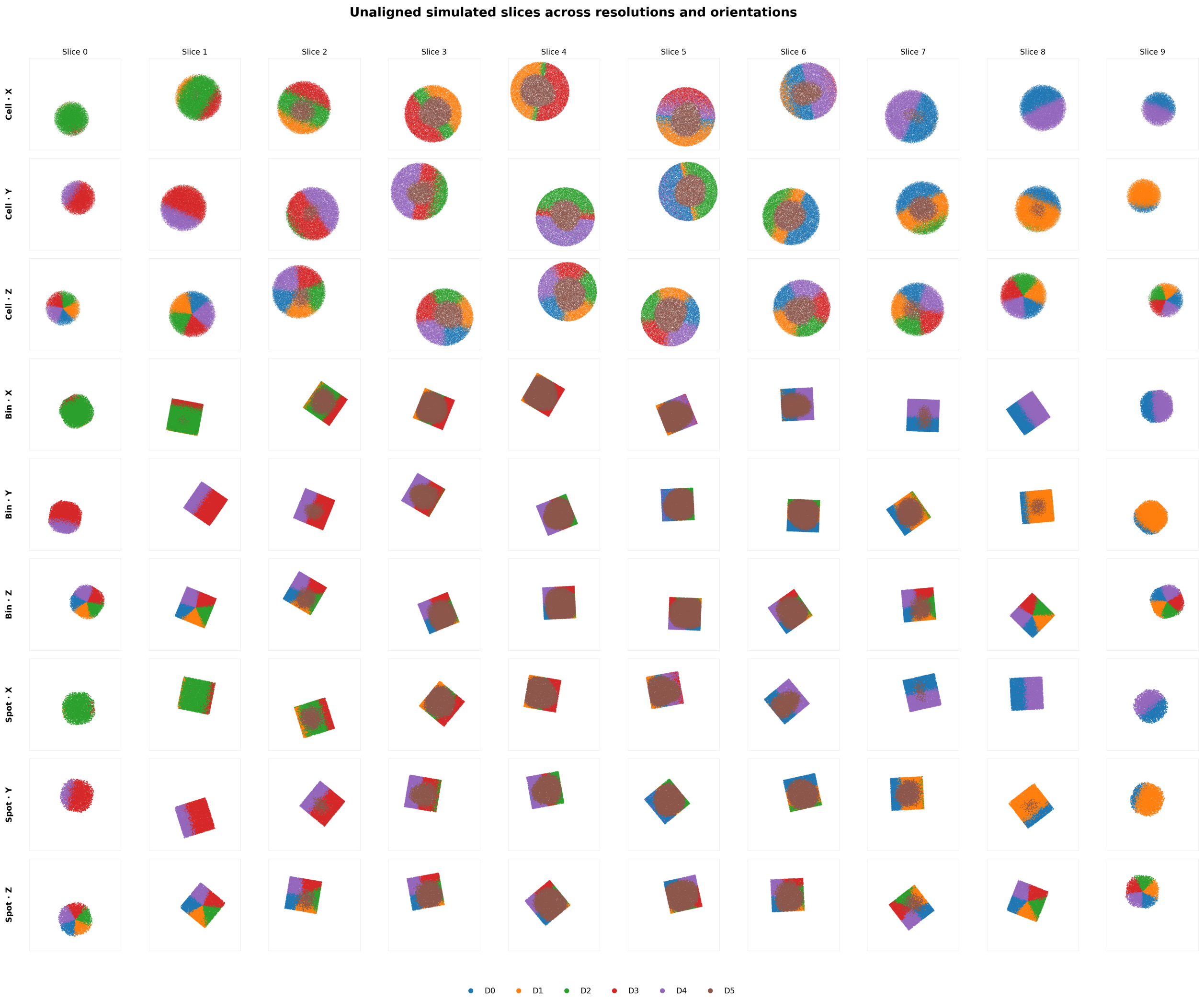


**Supplementary Figure S5: Unaligned simulated sections across three resolutions and three sectioning orientations.** Rows show cell-, bin-, and spot-level observations, each displayed for X, Y, and Z orientations. Observations are shown at their unaligned coordinates and colored by spatial domain.

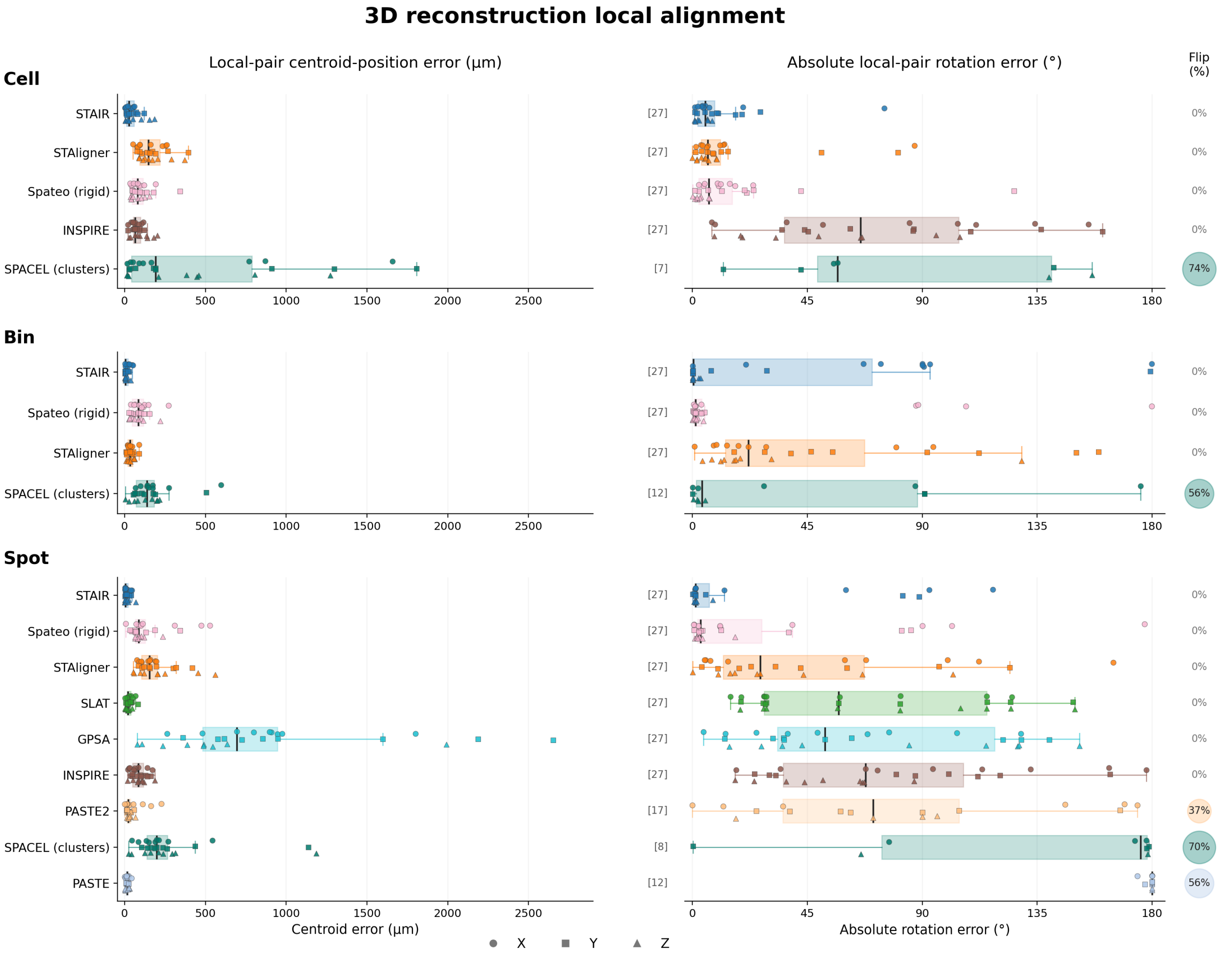


**Supplementary Figure S6: Local alignment errors and relative flips in simulated three-dimensional reconstructions.** Rows correspond to cell, bin, and spot resolutions. The first two columns show centroid-position errors and absolute rotation errors for consecutive section pairs, using the preceding section as the local reference. Points represent individual pairs pooled across X, Y, and Z orientations. Boxes indicate the interquartile range, center lines indicate medians, and whiskers extend to the most extreme observations within 1.5 times the interquartile range. Reflected pairs are excluded from rotation-error distributions. Circles beside the rotation plots summarize flip percentages among all 27 consecutive pairs per method and resolution.

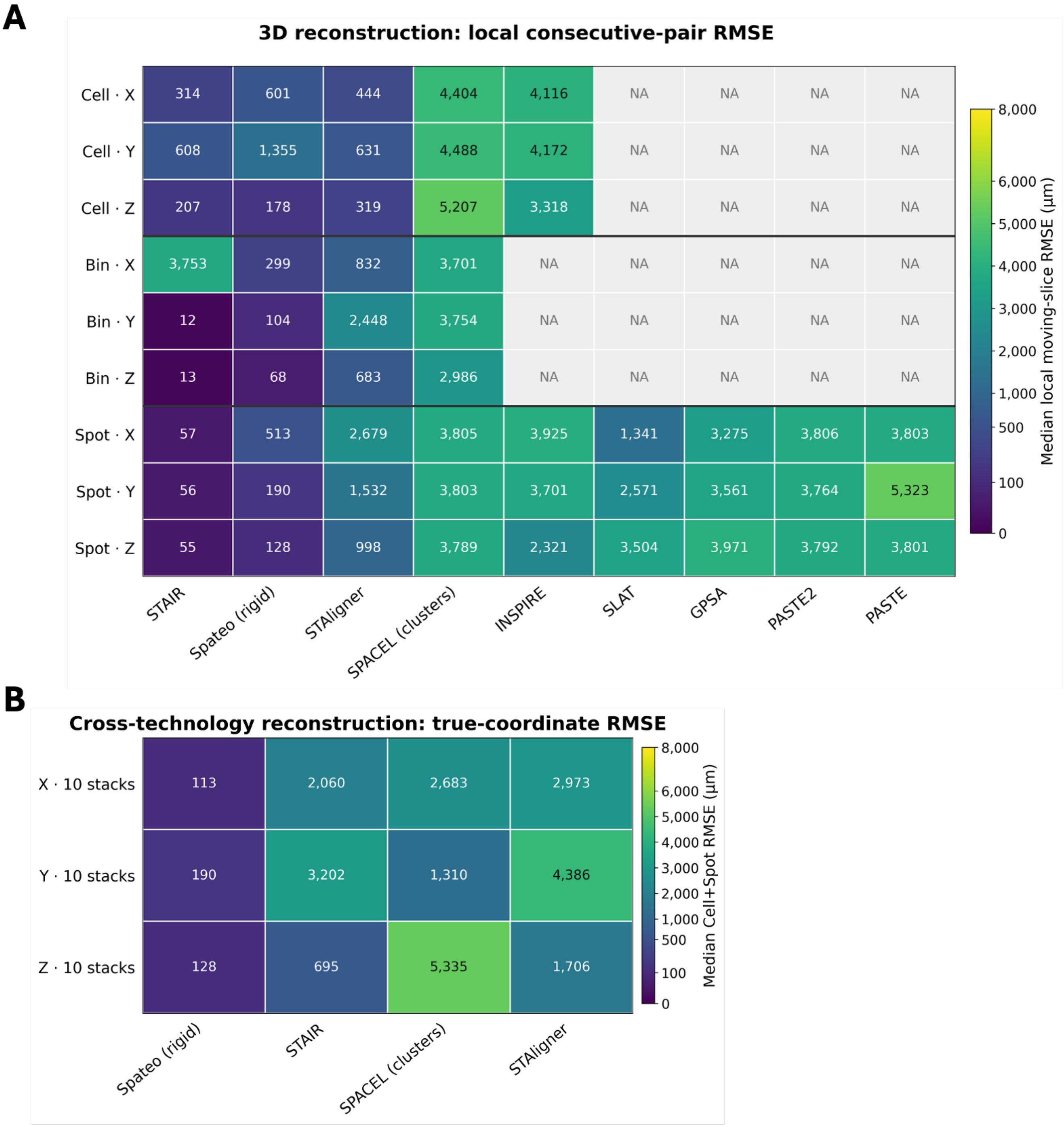


**Supplementary Figure S7: Coordinate-based reconstruction errors across simulated resolutions and sectioning orientations.** **(A)** Median local consecutive-pair root mean squared error (RMSE) for 3D reconstruction at cell, bin, and spot resolutions across X, Y, and Z orientations, using the preceding section as the local reference. **(B)** Median RMSE for cross-technology reconstruction across ten stacks per orientation, summarizing cell- and spot-level positional errors. Columns represent alignment methods. Heatmap colors and numerical values indicate RMSE in micrometers; NA denotes unavailable results.

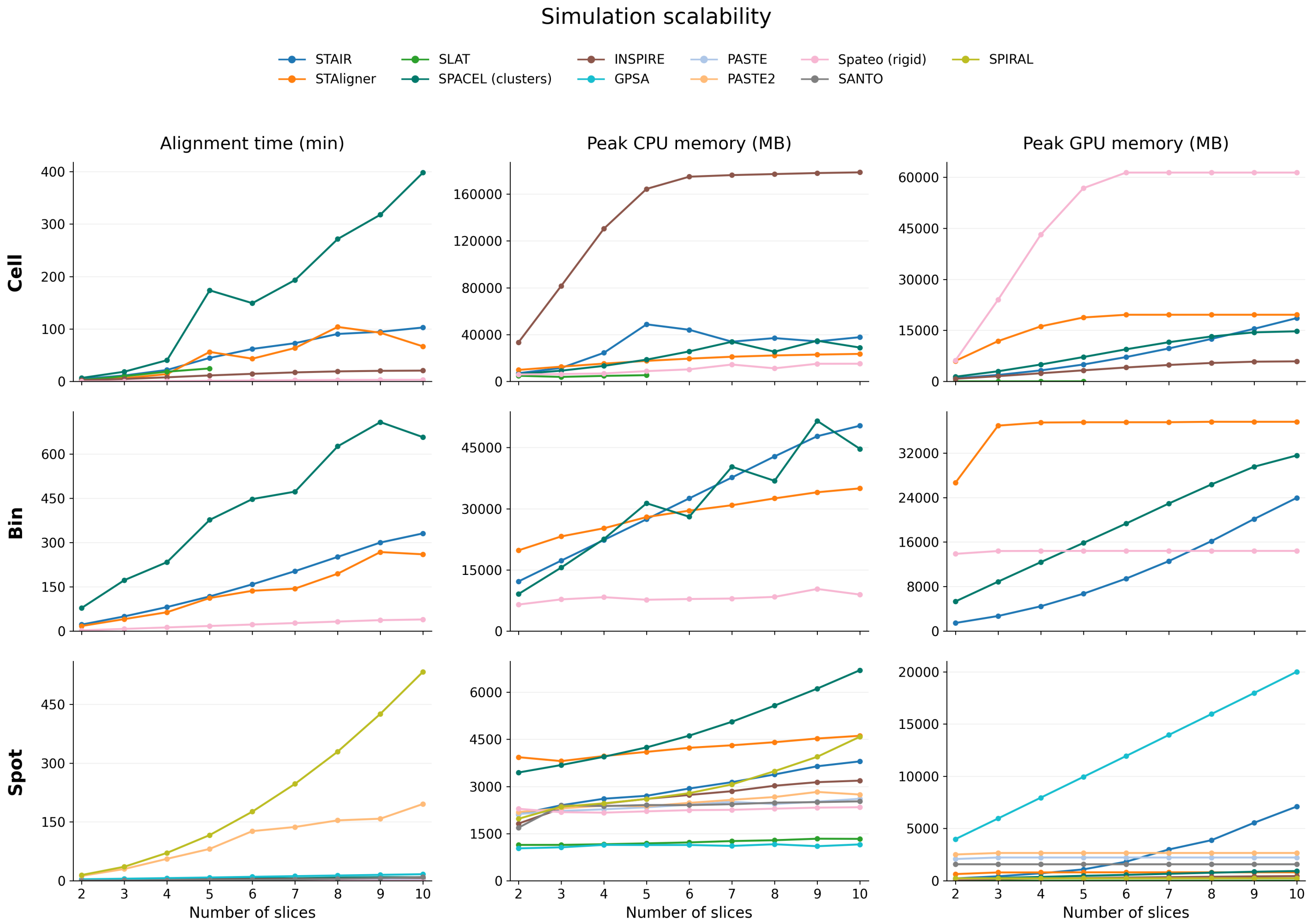


**Supplementary Figure S8: Computational scalability of alignment methods across simulated resolutions.** Rows correspond to cell, bin, and spot resolutions. Columns show alignment time, peak CPU memory, and peak GPU memory as the number of input sections increases from two to ten. Points represent median values across available sectioning orientations.

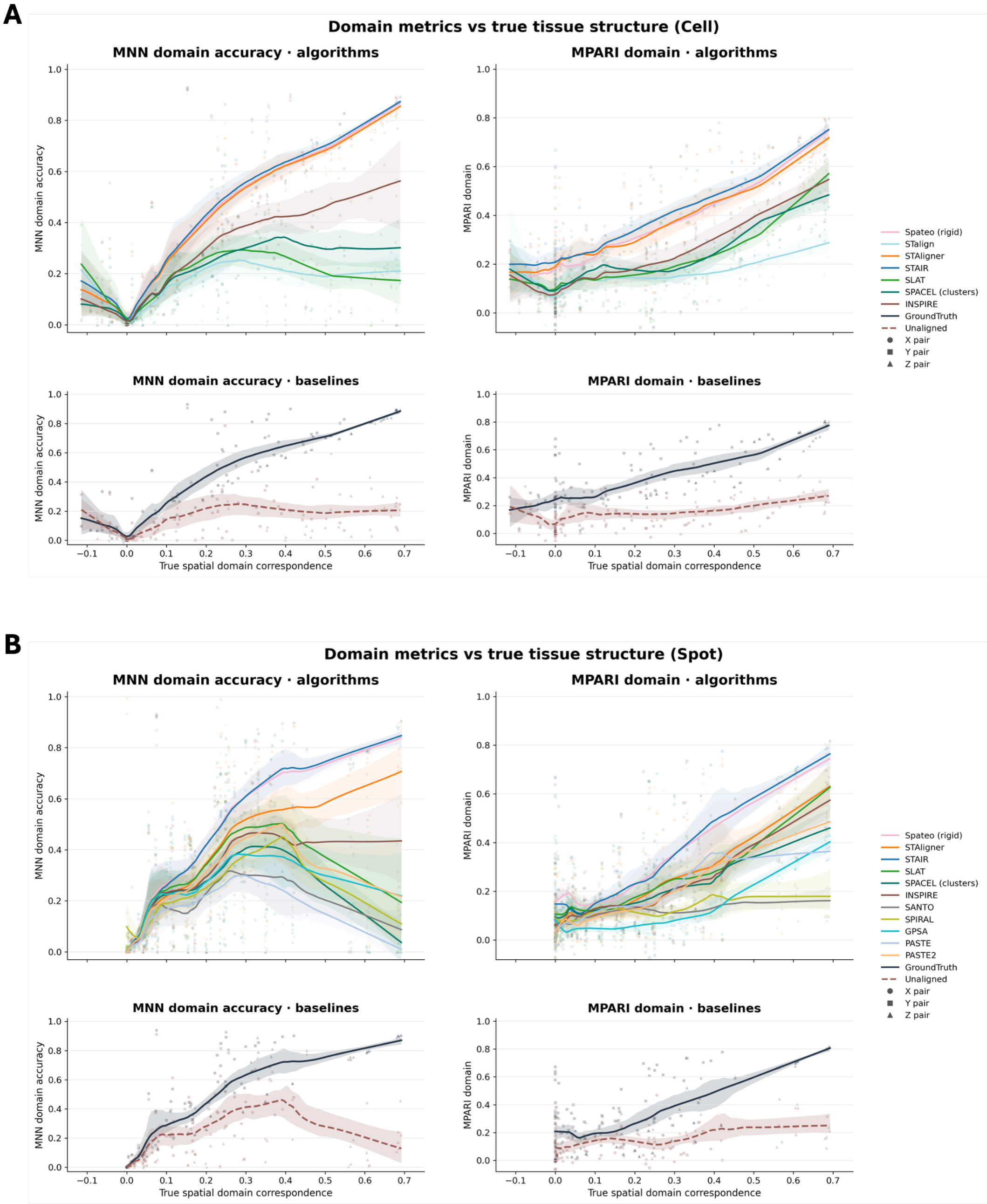


**Supplementary Figure S9: Domain-based alignment metrics versus true spatial domain correspondence in simulated data.** **(A)** and **(B)** show cell- and spot-level results, respectively. Within each block, columns show MNN domain accuracy and domain MPARI; the top row displays alignment methods, and the bottom row displays ground truth and unaligned baselines. Points represent individual section pairs, with marker shapes distinguishing X, Y, and Z orientations. Lines show smoothed trends, and shaded bands indicate the 2.5th–97.5th percentile range of pair-bootstrap fitted curves.

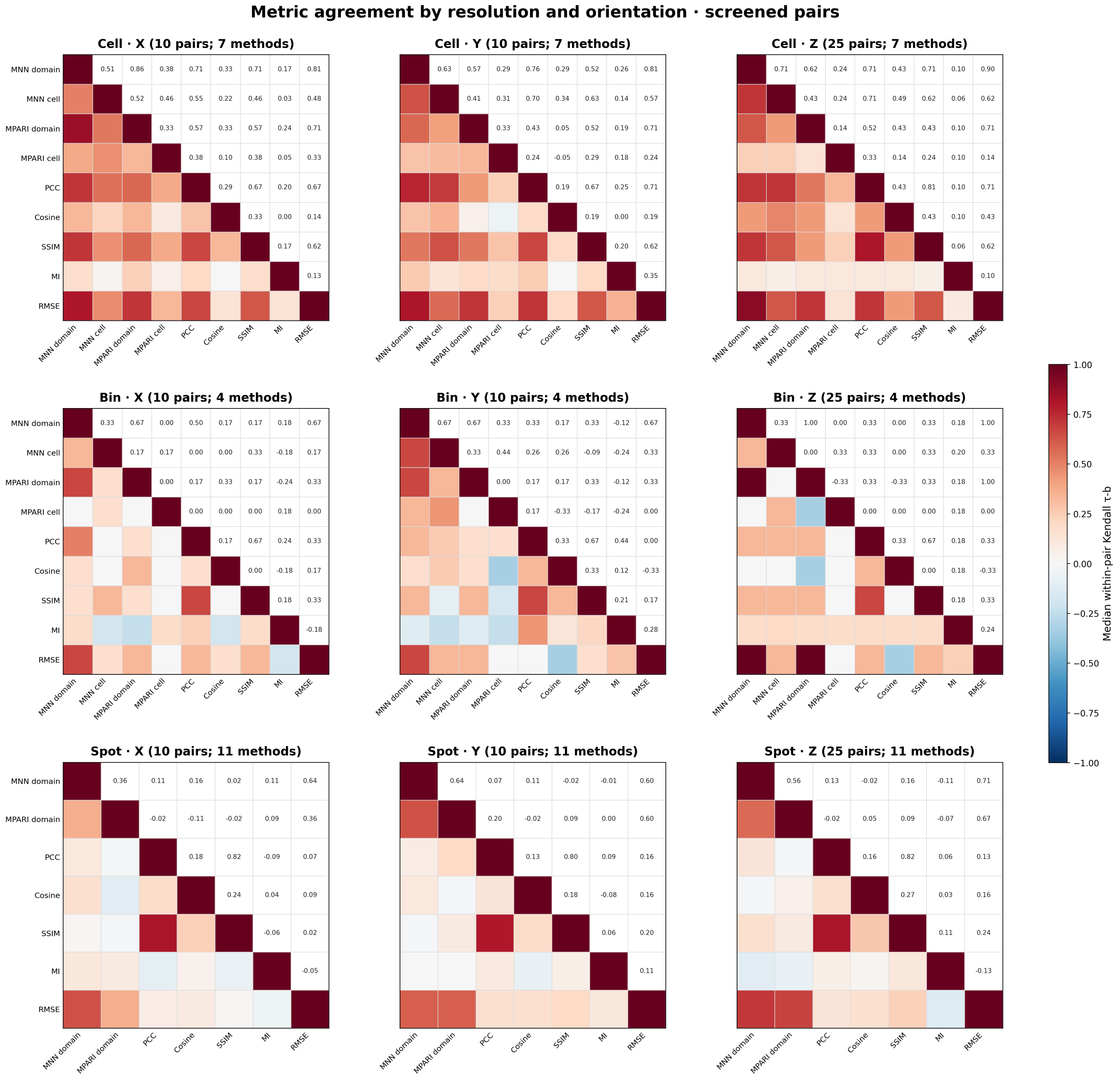


**Supplementary Figure S10: Metric ranking agreement across simulated resolutions and sectioning orientations.** Rows correspond to cell, bin, and spot resolutions, and columns correspond to X, Y, and Z orientations. Each heatmap summarizes agreement between metric-based algorithm rankings for screened section pairs. Colors in the lower triangle and numerical values in the upper triangle indicate Kendall's tau-b agreement. Metrics include domain- and cell-type-based MNN and MPARI, gene-expression-based PCC, cosine similarity, SSIM, and MI, and coordinate-based RMSE. Cell-type-based MNN and MPARI are omitted from spot-level panels.

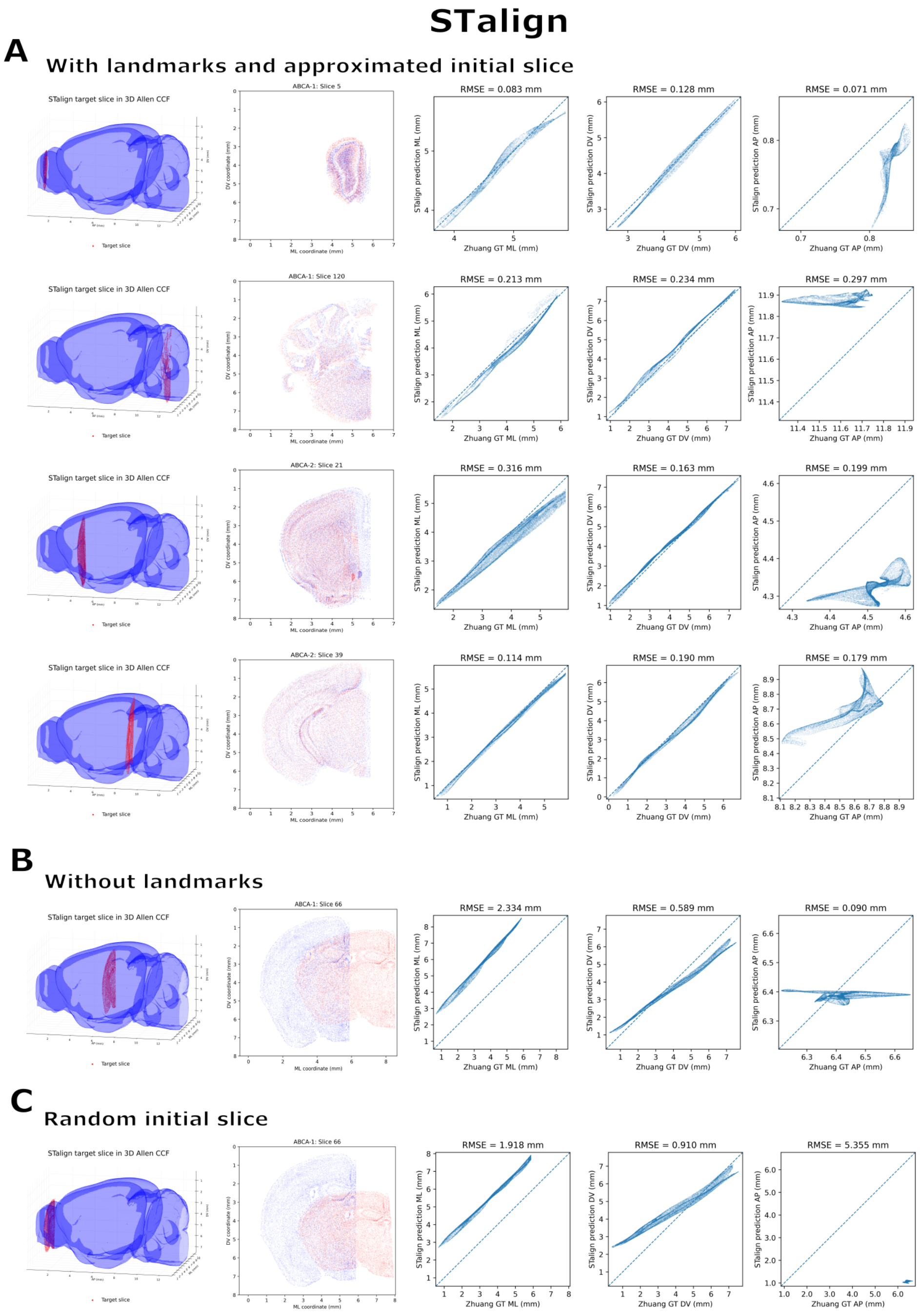


**Supplementary Figure S11: STalign registration across query sections and initialization settings.** (**A**) Registration of additional query sections from Zhuang-ABCA-1 and Zhuang-ABCA-2 using landmarks and an approximate initial atlas slice. (**B**) Registration of ABCA-1 slice 66 without landmarks. (**C**) Registration of the same query section without landmarks and with a randomly selected initial atlas slice. Each row shows the predicted section within the three-dimensional Allen CCF, an overlay of predicted and ground-truth coordinates in the mediolateral–dorsoventral plane, and predicted versus ground-truth coordinates along the mediolateral (ML), dorsoventral (DV), and anteroposterior (AP) axes. Dashed lines indicate perfect agreement ($y = x$), and axis-specific root mean squared errors (RMSE) are reported in millimeters.

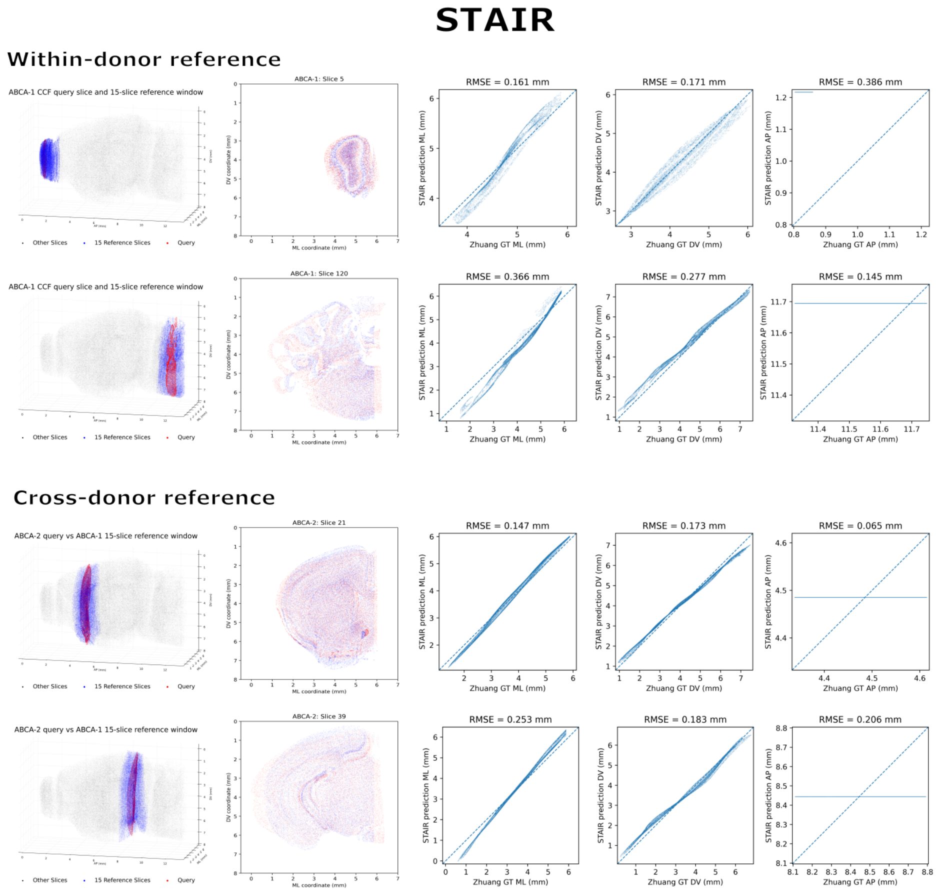


**Supplementary Figure S12: STAIR registration using within-donor and cross-donor reference sections.** Top: registration of Zhuang-ABCA-1 query slices 5 and 120 using reference sections from the same donor. Bottom: registration of Zhuang-ABCA-2 query slices 21 and 39 using reference sections from Zhuang-ABCA-1. Each row shows the query section within a three-dimensional 15-slice reference window, an overlay of predicted and ground-truth coordinates in the mediolateral–dorsoventral plane, and predicted versus ground-truth coordinates along the mediolateral (ML), dorsoventral (DV), and anteroposterior (AP) axes. Ground-truth coordinates correspond to the respective query donor. Dashed lines indicate perfect agreement ($y = x$), and axis-specific root mean squared errors (RMSE) are reported in millimeters.
